# Antibody design across observed and forecasted antigenic evolution

**DOI:** 10.64898/2026.11.10.765734

**Authors:** Soyoon Park, Talal Widatalla, Claudia L. Driscoll, Luis S. Mille-Fragoso, Ivan Specht, Xiaojing Gao, Brian L. Hie

## Abstract

The therapeutic efficacy of antibodies is often limited by the evolution of their targets. Designing or eliciting antibodies that withstand such evolution is challenging as their binding interfaces must accommodate antigenic variation. Here, we introduce Germinal Multi, a computational framework for *de novo* antibody design and rescue that jointly optimizes a single paratope against multiple antigenic variants. We applied Germinal Multi to cancer and viral antigens, targeting both observed antigenic variation and prospective antibody escape. Multi-target optimization yielded antibodies that bound resistance-associated EGFR variants or that neutralized antigenically distinct historical SARS-CoV-2 variants, with recognition extending to variants withheld from the design process. Beyond acting on observed evolution, we redesigned the broadly neutralizing antibody BD55-1205 to restore neutralization of a prospective escape variant engineered with guidance from an evolutionary forecasting model, while preserving activity against the parental variant. Successful candidates were identified after screening tens of designs per campaign. Together, these results establish multi-target computational design as a strategy for generating antibody breadth across observed antigenic variation and demonstrate the potential of evolutionary forecasting to guide antibody redesign against prospective escape.

## 1. Introduction

The ability of antibodies to recognize molecular targets with high specificity has enabled their therapeutic use across infectious disease, cancer, and other areas of medicine. However, antigenic evolution can limit their effectiveness, as mutations that impair antibody recognition emerge over time and reduce therapeutic efficacy. For example, successive waves of SARS-CoV-2 evolution rendered several therapeutic monoclonal antibodies targeting the receptor-binding domain (RBD) ineffective against emerging variants, prompting restrictions on their use less than two years after their initial authorization (Starr et al., 2021; Zitek et al., 2023; Alcantara et al., 2023).

Antigenic evolution has motivated efforts to develop broadly neutralizing antibodies (bnAbs) that recognize conserved features of their targets and retain activity across antigenic variants. bnAbs have been identified through antibody isolation from convalescent or vaccinated individuals (Cao et al., 2022a), animal immunization (Zhou et al., 2017), and *in-vitro* screening of large-scale antibody libraries (Sui et al., 2009), all of which can be resource intensive. Immunization strategies, including multivalent vaccination, immunofocusing, and germline targeting, aim to elicit broader antibody responses by directing immunity toward conserved epitopes or recruiting appropriate B-cell lineages (Cohen et al., 2022; Yassine et al., 2015; Leggat et al., 2022). However, these strategies can require substantial immunogen engineering, and achieving the desired breadth depends on recruiting and guiding the maturation of appropriate B-cell lineages, which can be challenging (Yassine et al., 2015; Leggat et al., 2022). Moreover, even antibodies with broad activity against existing antigenic variants may ultimately lose activity against future variants (Cao et al., 2022b).

These challenges motivate computational approaches that can directly design antibodies with the desired breadth. Recent advances in *de novo* antibody design have produced experimentally validated, epitope-specific antibodies (Mille-Fragoso et al., 2026; Bennett et al., 2025). For example, Germinal identified binding antibodies to four targets after experimentally testing 43–101 designs per target, illustrating the potential to reduce reliance on animal immunization and large-scale experimental library screening (Mille-Fragoso et al., 2026). However, these *de novo* methods were demonstrated by designing against a single antigen structure at a time, without simultaneously optimizing a single paratope against multiple variants of a shared epitope.

Here, we extend computational antibody design to generate antibodies with broad binding or neutralization activity across antigenic variants. We build on our previous single-target antibody design pipeline, Germinal (Mille-Fragoso et al., 2026), by formulating antibody design against multiple antigenic variants as a multi-objective optimization problem. The resulting framework, Germinal Multi, jointly optimizes structural confidence in predicted complexes between a single antibody paratope and multiple target variants. Germinal allows explicit targeting of conserved structural regions, while simultaneous modeling of multiple variants accounts for substitutions within the binding interface.

We applied this framework to cancer and viral disease systems in which antigenic variation contributes to clinically observed antibody escape. In cancer, we selected epidermal growth factor receptor (EGFR), an oncogenic receptor tyrosine kinase for which acquired resistance to the approved antibody cetuximab can arise through substitutions in the antigen itself (**Figure 1A**). In viral infectious disease, we selected the SARS-CoV-2 spike receptor-binding domain (RBD), against which therapeutic antibodies have repeatedly lost activity following viral evolution. We jointly optimized antibodies against the antigenically distinct BA.2 and KP.3 variants to generate candidates that neutralize both (**Figure 1B**). Across these settings, Germinal Multi yielded antibodies with broad recognition across antigenic variants after experimental screening of tens of designs per campaign, comparable to the low-*n* screening scale of single-target Germinal campaigns (Mille-Fragoso et al., 2026) (**Figure 1D**). To investigate potential future viral escape in a currently circulating SARS-CoV-2 lineage, XFG, we integrated predictions from antiGen, our computational method for forecasting viral evolution (Specht et al., 2026), with deep mutational scanning data obtained in a KP.3.1.1 background (Dadonaite et al., 2025). We then experimentally evaluated whether substitutions forecasted by antiGen for XFG at sites associated with BD55-1205 escape in KP.3.1.1 also conferred escape in the XFG background (**Figure 1C**). Using Germinal Multi, we redesigned BD55-1205 to restore neutralization of an engineered XFG escape variant while preserving activity against parental XFG. These results support the use of evolutionary forecasting to guide antibody redesign, offering a path toward evolutionarily resilient therapeutics.

**Figure 1.**
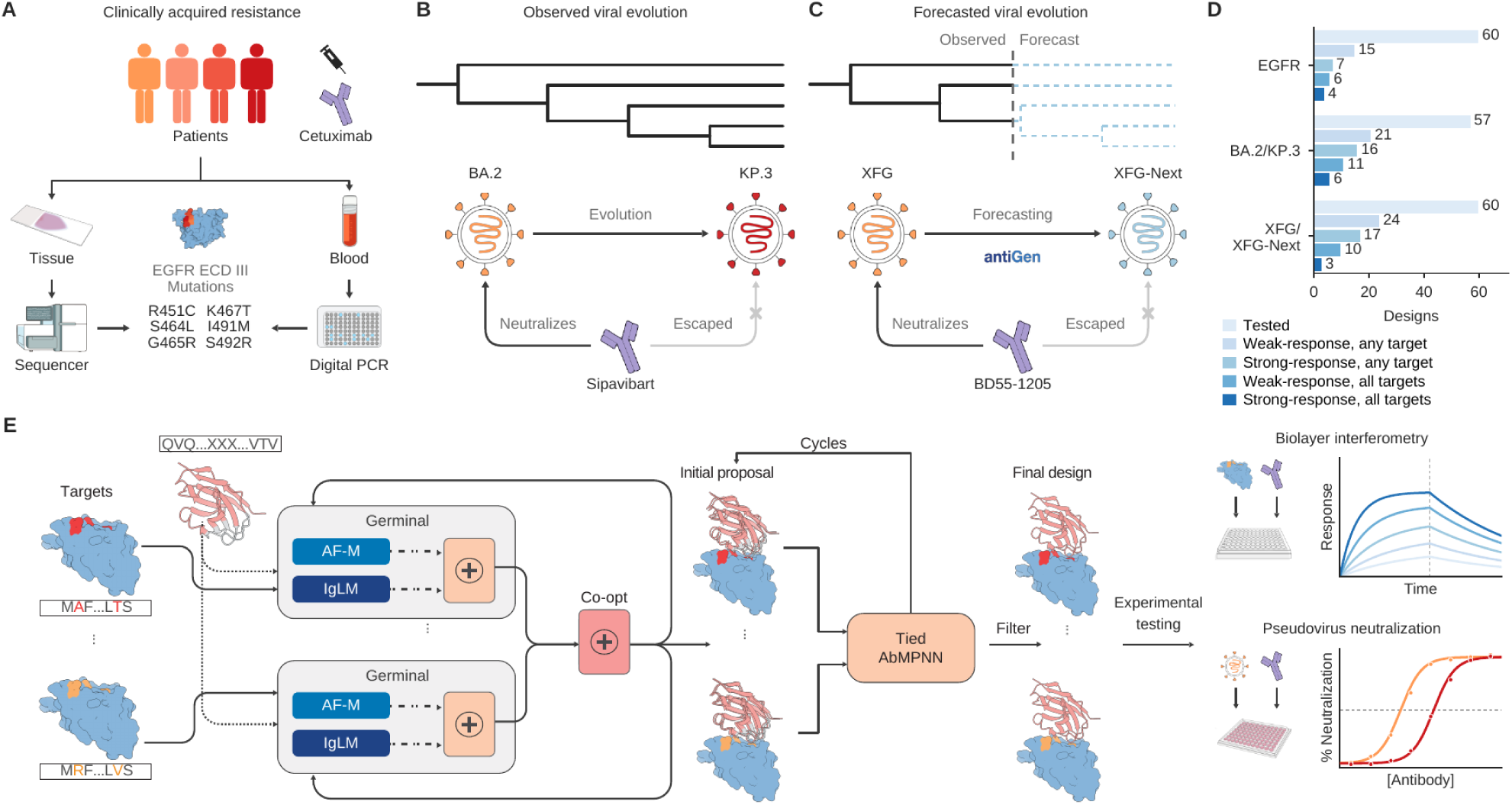
Germinal Multi jointly optimizes antibody design across observed and forecasted antigenic variation. **(A)** Resistance-associated variation in EGFR motivates antibody design across multiple antigen variants. The schematic illustrates tissue sequencing and digital PCR of blood samples as approaches for detecting resistance-associated substitutions. The selected panel comprises six substitutions in EGFR domain III supported by previous studies describing clinical or preclinical data: R451C, S464L, G465R, K467T, I491M, and S492R (Arena et al., 2015; Sánchez-Martín et al., 2016; Montagut et al., 2012). **(B)** Observed viral evolution motivates joint optimization against antigenically distinct SARS-CoV-2 variants. AZD3152 (Sipavibart) neutralizes BA.2 but loses activity against KP.3. The tree illustrates diversification schematically and does not represent an inferred phylogeny. **(C)** Forecast-informed antibody redesign against a prospective challenge antigen. Predictions from antiGen and published deep mutational scanning measurements obtained in a SARS-CoV-2 KP.3.1.1 background informed the selection of substitutions subsequently evaluated in the XFG background. The resulting engineered challenge variant, XFG-Next, escapes neutralization by parental BD55-1205. Solid black branches represent observed evolution; dashed blue branches represent forecasted evolution possibilities. **(D)** BLI screening outcomes for the three antibody-design campaigns. For a given target, categories are not mutually exclusive. *Tested* denotes the number of designs experimentally evaluated. Weak-response and strong-response categories use reference-subtracted BLI response thresholds of 0.05 and 0.5 nm, respectively, measured at the end of the association phase. “Any target” denotes meeting the corresponding threshold against at least one antigen; “all targets” denotes meeting it against every antigen in the experimental panel. Categories overlap. The experimental antigen panels comprise wild-type EGFR-Fc and six individual resistance variants, BA.2 and KP.3 RBDs, or XFG and XFG-Next RBDs. For EGFR, the six designs exceeding the strong-response threshold against wild-type EGFR were subsequently screened against all six individual resistance variants; counts for responses against all targets therefore represent lower bounds. **(E)** Germinal Multi workflow. A shared antibody sequence is evaluated against multiple antigen targets. AlphaFold-Multimer (AF-M) structure-based terms and IgLM pseudo-likelihood contribute to joint optimization (Shuai et al., 2023). Initial antibody–antigen proposals undergo iterative sequence-design cycles using tied AbMPNN, which conditions on multiple structures while enforcing a single sampled antibody sequence. Designs are filtered and evaluated experimentally by BLI and, for viral targets, pseudovirus neutralization assays. Red and orange antigen regions indicate sequence differences between antigen targets.

## 2. Results

### 2.1. Extending Germinal to multi-epitope co-optimization

We extended our previous method, Germinal (Mille-Fragoso et al., 2026), to support antibody design against multiple targets, yielding Germinal Multi (**Figure 1E**). In addition to the *de novo* pipeline, which freely designs the complementarity-determining regions (CDRs), we introduce a constrained redesign pipeline that biases designs toward the parental antibody’s CDR sequences and a reference antibody-antigen binding pose. This “rescue” configuration is intended to restore activity to existing therapeutic antibodies affected by antigenic evolution while retaining desirable properties, such as developability and binding to the ancestral antigen (**Methods**).

Alongside improvements to computational efficiency (**Methods**), Germinal Multi introduces three principal features for co-optimization against multiple targets. First, during the design stage, we evaluate prediction confidence for the antibody in complex with each target and combine the corresponding gradients. Second, during the sequence design stage, instead of designing CDR residues conditioned on a single target, we use a multi-target sequence design approach with AbMPNN (Dreyer et al., 2023), sampling from the normalized geometric mean of the antibody sequence conditioned on each target. Finally, at the filtering stage, we evaluate all filters to each target, selecting designs with confident predictions and similar predicted binding modes across all targets. Across multiple campaigns, Germinal Multi co-optimization improves multi-target binding relative to single-target optimization with Germinal, as assessed *in silico* (**Figures S29, S30 and S31)**. Further details on the co-optimization pipeline are provided in **Methods**.

### 2.2. Designing broad recognition of clinically observed cetuximab-resistance mutations to EGFR

Oncoproteins are common drug targets, and mutations in these proteins can confer resistance to therapeutic antibodies. EGFR, a receptor tyrosine kinase implicated in several cancers, is targeted by monoclonal antibodies (mAbs) such as cetuximab and panitumumab in colorectal cancer. Resistance to these anti-EGFR mAbs can arise through several mechanisms (Misale et al., 2012; Bardelli et al., 2013; Montagut et al., 2012). Among these mechanisms, missense mutations in the EGFR extracellular domain (ECD) that impair antibody binding or activity have been reported in approximately 15–20% of patients with colorectal cancer and acquired resistance to EGFR blockade (Sánchez-Martín et al., 2016). One strategy to address resistance mediated by EGFR ECD mutations is to combine antibodies targeting non-overlapping epitopes into a single therapeutic product (Pedersen et al., 2010; Sánchez-Martín et al., 2016; Arena et al., 2016). Motivated in part by the formulation and analytical complexity associated with antibody cocktails (Kim et al., 2020; Kuhne et al., 2023), we sought to design a single antibody that retains binding across diverse resistance-associated substitutions within or near its epitope.

Cetuximab blocks binding of endogenous EGFR ligands such as EGF or TGF-*α* and sterically hinders the adoption of the extended ECD conformation required for receptor dimerization and signaling activation (Li et al., 2005). To target this clinically-validated, functionally relevant region, we specified the cetuximab epitope on EGFR domain III as our target epitope. We selected six previously characterized substitutions in this domain associated with clinical or preclinical resistance to cetuximab or related anti-EGFR antibodies: R451C, S464L, G465R, K467T, I491M and S492R (Arena et al., 2015; Sánchez-Martín et al., 2016; Montagut et al., 2012) (**Figure 2A**). These substitutions were identified in patients treated with anti-EGFR antibodies or in cetuximab-resistant cell models. The corresponding variants exhibit distinct antibody-sensitivity profiles, ranging from preferential impairment of cetuximab recognition to cross-resistance to both cetuximab and panitumumab. Five substitutions lie within the cetuximab-binding surface, whereas R451C lies outside the direct epitope (Arena et al., 2015; Sánchez-Martín et al., 2016).

**Figure 2.**
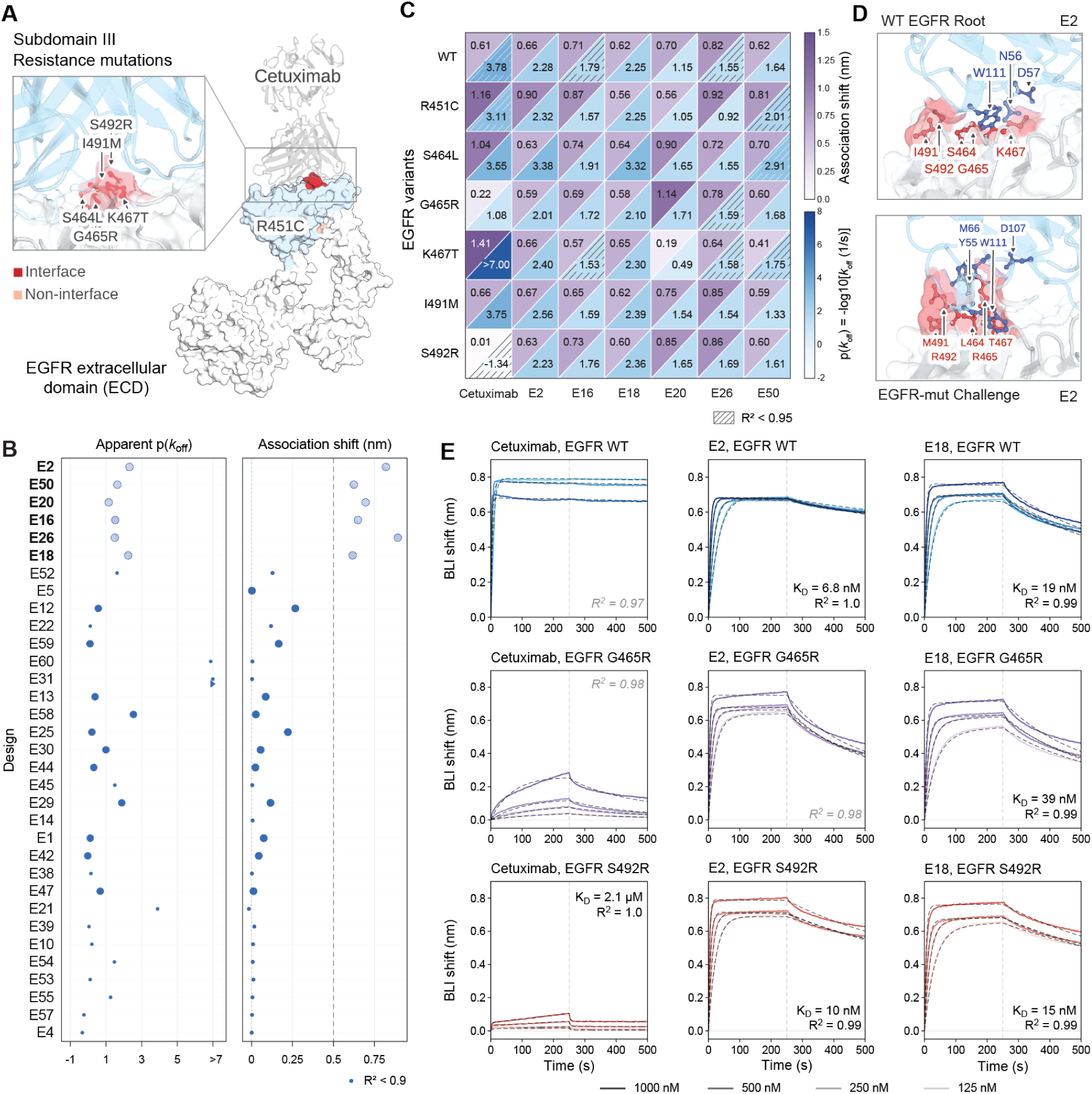
Multi-epitope antibody design enables binding across resistance-associated EGFR variants. **(A)** Structural view of cetuximab bound to the EGFR extracellular domain (ECD), with an enlarged view of resistance-associated positions in domain III. Positions within and outside the cetuximab interface are highlighted in red and peach, respectively. **(B)** Single-concentration BLI screening of antibody designs against wild-type EGFR. Designs were generated by Germinal Multi using a trastuzumab framework by jointly optimizing binding to wild-type EGFR and a composite challenge antigen containing all six selected resistance-associated substitutions. Plots show association responses and dissociation rate constant estimates expressed as *p*(*k*_off_) = − log_10_ [*k*_off_/(s^−1^)]; higher values indicate slower dissociation. Smaller symbols denote kinetic fits with *R*^2^ < 0.9. The dashed line marks an association response of 0.5 nm, and hatched symbols identify designs exceeding this threshold. Bold labels identify designs shown in **(C)** Binding profiles of cetuximab and designs E2, E16, E18, E20, E26, and E50 against wild-type EGFR and the six individual resistance-associated variants. Within each cell, the upper-left triangle shows association responses in nanometers, and the lower-right triangle shows the apparent *p*(*k*_off_) estimate. Hatching indicates kinetic fits with *R*^2^ < 0.95. In **(B,C)**, antibodies were tested at 1,000 nM; values labeled > 7 indicate *k*_off_ < 10^−7^ s^−1^. **(D)** Structural models of E2 bound to wild-type EGFR and the composite challenge antigen. Selected antibody residues are labeled in blue; antigen residues at resistance-associated positions are labeled in red. **(E)** BLI titrations of cetuximab, E2, and E18 (columns, left to right) against wild-type EGFR (top row, blue), G465R (middle row, purple), and S492R (bottom row, red). Antibody concentrations were 1,000, 500, 250, and 125 nM. Colored solid traces show reference-subtracted responses; black dashed traces show kinetic fits. Vertical dotted lines mark the start of dissociation at 250 s. The same y-axis limits are used across antibodies within each target. Apparent *K_D_* estimates and corresponding *R*^2^ values are annotated for fits with *R*^2^ ≥ 0.99; otherwise, only *R*^2^ values are shown in gray italics. Kinetic fitting is described in **Methods**.

During computational design, rather than including all six individual EGFR variants as separate optimization targets, we co-optimized antibodies against wild-type EGFR and an artificial composite antigen containing all six resistance-associated substitutions, termed the “challenge” antigen. We hypothesized that this strategy would promote recognition across the individual variants and potentially combinations thereof, while reducing the number of optimization targets and associated computational cost. We redesigned all six CDRs within a trastuzumab framework, which we previously determined to be amenable to CDR alteration (Mille-Fragoso et al., 2026), and generated 60 candidate sequences using our co-optimization pipeline (**Methods**).

We evaluated all 60 designs for binding to wild-type EGFR by single-concentration biolayer interferometry (BLI) at an antibody concentration of 1, 000 nM. To prioritize candidates for further characterization, we defined an initial hit as an antibody with a response exceeding 0.5 nm at the end of the association phase against wild-type EGFR (**Figure 2B**; **Figures S1 and S2**). Six designs—E2, E16, E18, E20, E26, and E50—met this threshold and were subsequently screened against all six individual resistance-associated EGFR variants at the same single antibody concentration (**Figure 2C**) and further characterized by BLI titration.

Four designs—E2, E16, E18, and E26—exceeded the 0.5 nm response threshold against wild-type EGFR and all six individual resistance-associated variants (**Figure 2C**). E50 showed similarly broad recognition, with responses exceeding 0.5 nm against six targets and a response of 0.41 nm against K467T. Notably, multiple designs retained binding to both G465R and S492R, while cetuximab exhibited markedly reduced binding under the same assay conditions (**Figure 2C, E**; **Figures S3 and S4**). Structural models from AlphaFold 3 (Abramson et al., 2024) of representative design E2 bound to wild-type EGFR and the composite challenge antigen illustrate the predicted interactions at resistance-associated positions (**Figure 2D**).

These results demonstrate that co-optimization against wild-type EGFR and a composite challenge antigen can yield antibodies that recognize a therapeutically relevant antigen and retain binding across multiple individually tested EGFR resistance-associated variants.

### 2.3. Designing antibody breadth across historical SARS-CoV-2 evolution

Beyond cancer, we applied Germinal Multi to problems in viral evolution, where successive waves of antigenic evolution have repeatedly eroded the activity of therapeutic antibodies (Planas et al., 2022; Cao et al., 2023). We focused on the epitope of the SARS-CoV-2 spike RBD recognized by AZD3152 (sipavibart), a clinical antibody that retained neutralizing activity against antigenically divergent Omicron variants, including BA.2 and JN.1, but lost neutralizing activity against KP.3 (Planas et al., 2024; Cai et al., 2024) (**Figure 3A, C**). Corresponding BLI measurements are shown in **Figure S6E**.

**Figure 3.**
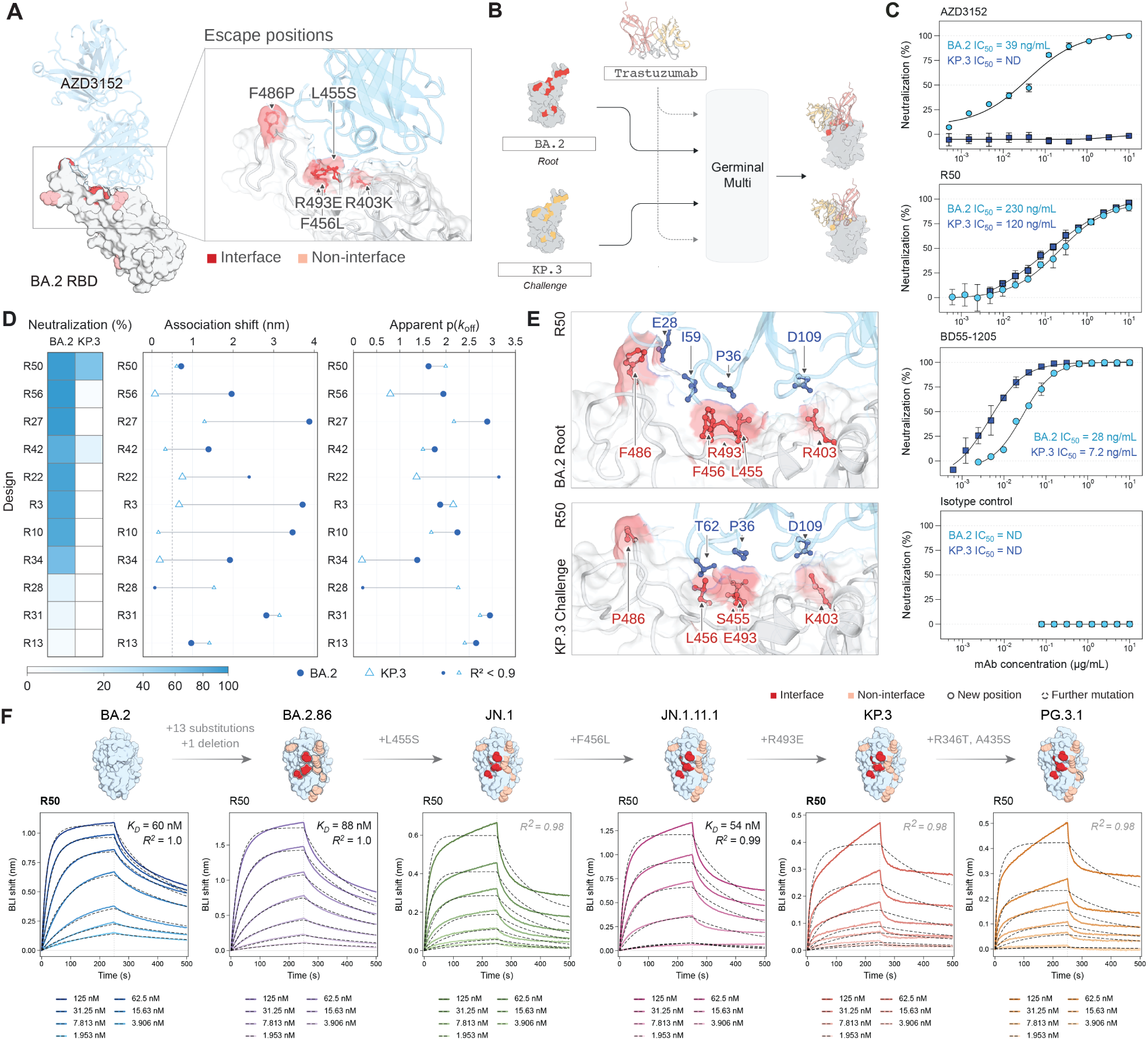
Antibody design spanning historical SARS-CoV-2 evolution. **(A)** Structural view of AZD3152 bound to the BA.2 receptor-binding domain (RBD), with an enlarged view of selected positions that differ between BA.2 and KP.3. Positions within and outside the antibody–RBD interface are highlighted in red and peach, respectively. **(B)** Schematic of Germinal Multi co-optimization against BA.2 and KP.3 using a trastuzumab framework. **(C)** Neutralization of BA.2 and KP.3 pseudoviruses by R50, parental reference antibody AZD3152, positive control BD55-1205, and the 13D4 isotype control. Two independent experiments were performed, each with two technical replicates per concentration. Points show the mean of the two experiment-level means; error bars indicate SD (*n* = 2 independent experiments). Lines show neutralization curves fitted as described in **Methods**. Annotated IC_50_ values are the geometric means of the unrounded fitted estimates from two independent experiments, expressed in ng/mL. Experiment-specific fits are shown in **Figure S11**. ND indicates that an IC_50_ could not be determined within the tested concentration range. **(D)** Single-concentration neutralization and BLI screening results for the top 11 of 57 screened antibody designs, ranked by mean percent neutralization across BA.2 and KP.3 at 10 *μ*g/mL. Heatmap shows mean percent neutralization from two independent experiments, each performed in technical duplicate. Adjacent plots show BLI association responses and apparent dissociation rates expressed as *p*(*k*_off_) = − log_10_ [*k*_off_/(s^−1^)], measured at 1,000 nM antibody; higher values indicate slower dissociation. Filled circles represent BA.2; open triangles represent KP.3. Lines connect measurements for the same design; smaller symbols indicate kinetic fits with *R*^2^ < 0.9. Full sensorgrams and additional screening results are shown in **Figures S8, S9, and S10A**. Neutralization titrations for designs classified as BA.2-only neutralizers in the initial screen are shown in **Figure S13**. **(E)** Structural models of R50 bound to BA.2 and KP.3. Selected antibody residues are labeled in blue; antigen residues at positions differing between the two variants are labeled in red. **(F)** R50 binding titrations against BA.2, BA.2.86, JN.1, JN.1.11.1, KP.3, and PG.3.1 RBDs. Colored solid traces show reference-subtracted responses at the indicated antibody concentrations; black dashed traces show kinetic fits. Vertical dotted lines mark the start of dissociation at 250 s. Apparent *K_D_* estimates and corresponding *R*^2^ values are annotated for fits with *R*^2^ ≥ 0.99; otherwise, only *R*^2^ values are shown in gray italics. Structural views above the sensorgrams illustrate substitutions across the variant series. Positions within and outside the antibody–RBD interface are highlighted in red and peach, respectively. Solid outlines mark newly altered positions; dashed outlines mark further substitutions at previously altered positions. Only BA.2 and KP.3 were included during co-optimization. Kinetic fitting is described in **Methods**.

To test whether computational antibody design could generate antibodies active against both an earlier variant and a variant that escapes the reference antibody, we used Germinal Multi for *de novo* antibody design targeting the AZD3152 epitope on BA.2 and KP.3, jointly optimizing all six complementarity-determining regions within a trastuzumab framework (**Figure 3B**; **Methods**). We hypothesized that simultaneous optimization against these two antigenically distinct targets could yield antibodies that bind and neutralize both variants, with recognition extending to evolutionary intermediates not explicitly included during design.

We screened 57 Germinal Multi designs for binding and neutralization against BA.2 and KP.3. The top 11 designs, ranked by mean percent neutralization across the two variants, are shown in **Figure 3D**. Combined screening identified R50 as a lead with activity against both variants (**Figures S8, S9 and S10A**). Pseudovirus neutralization titrations yielded IC_50_ values of 230 ng/mL against BA.2 and 120 ng/mL against KP.3 (**Figure 3C**; **Figure S11**). Structural models illustrate the predicted interactions of R50 with BA.2 and KP.3 (**Figure 3E**). BLI binding titrations of R50 yielded apparent *K_D_* estimates of 60 nM against BA.2 and 20 nM against KP.3 (**Figure 3F**).

R31 showed the strongest combined single-point BLI association responses across the two RBDs, but neutralized only BA.2 in the single-concentration screen (**Figure 3D**), with subsequent titrations yielding a geometric mean IC_50_ of 2.7 µg mL^−1^ against BA.2 and no determinable IC_50_ against KP.3. Despite more moderate BLI responses, R42 neutralized both variants, with geometric mean IC_50_ values of 0.48 and 3.2 µg mL^−1^ against BA.2 and KP.3, respectively (**Figure S12**). These results underscore the importance of evaluating neutralization alongside binding (**Figure S12**).

As an alternative to *de novo* design, we evaluated baseline approaches that introduced substitutions into AZD3152 using the structure-conditioned models ESM-IF1 and FAMPNN or amino-acid frequencies from human antibody sequences in abYsis (Shanker et al., 2024; Widatalla et al., 2025; Swindells et al., 2017) (**Figure S6A**). We screened individual substitutions (**Figures S5 and S6C**) followed by combinations of promising substitutions (**Figures S6B and S7**) for BA.2 and KP.3 binding by BLI (**Methods**). The selected pentuple mutant of AZD3152 showed modest recovery of KP.3 binding while retaining BA.2 binding, but failed to neutralize either KP.3 or BA.2 pseudoviruses (**Figure S6D**). In contrast, Germinal Multi generated an antibody that neutralized both variants after a single round of experimental screening.

We next asked whether R50’s recognition extended beyond the two antigen sequences used during optimization. Notably, R50 also bound BA.2.86, JN.1 and JN.1.11.1 RBDs, none of which were included during optimization, demonstrating binding breadth across evolutionary intermediates (**Figure 3F**; **Figure S14**). R50 additionally retained binding to the RBD of a subsequent variant, PG.3.1, with an apparent *K_D_* of 26 nM (*R*^2^ = 0.98) (**Figure 3F**). Together, these results demonstrate that Germinal Multi can generate an antibody that neutralizes the antigenically distinct historical SARS-CoV-2 variants BA.2 and KP.3, with binding breadth extending to evolutionary intermediates and a subsequent variant not included during optimization.

### 2.4. Designing antibody breadth across forecasted SARS-CoV-2 evolution

Finally, we investigated whether co-optimization, informed by evolutionary forecasting and experimental datasets, could support proactive antibody development. We focused on BD55-1205, a broadly neutralizing antibody with activity against the SARS-CoV-2 XFG lineage, which is the dominant circulating variant at the time of this study. To identify prospective antigenic variation, we used our recently developed model, antiGen, which uses the phylogenetic structure of past antigenic evolution to forecast substitutions (Specht et al., 2026).

We sought to identify XFG mutations that were (i) supported by evolutionary forecasting, (ii) compatible with viral entry, and (iii) associated with reduced susceptibility to BD55-1205 neutralization. To inform candidate prioritization, we consulted a published pseudovirus-based deep mutational scanning (DMS) landscape of BD55-1205 escape and ACE2 binding obtained in a KP.3.1.1 spike background (Dadonaite et al., 2025). Because corresponding XFG DMS data were unavailable, we used the KP.3.1.1 landscape to prioritize candidates for experimental assessment in the XFG background.

The published DMS landscape identified six major BD55-1205 escape-associated sites in the RBD (405, 456, 475, 493, 501, and 505). Predictions from the site-independent version of antiGen (Specht et al., 2026) included substitutions at three of these six sites: A475V, E493R, and Y501T (**Figure 4A,B**). Of these three predicted substitutions, A475V was subsequently detected in circulating SARS-CoV-2 sequences (**Figure S15**). All three single-substitution variants retained measurable pseudovirus infectivity, supporting compatibility with viral entry under the assay conditions, although E493R and Y501T showed lower infectivity than parental XFG (**Figure S16**).

**Figure 4.**
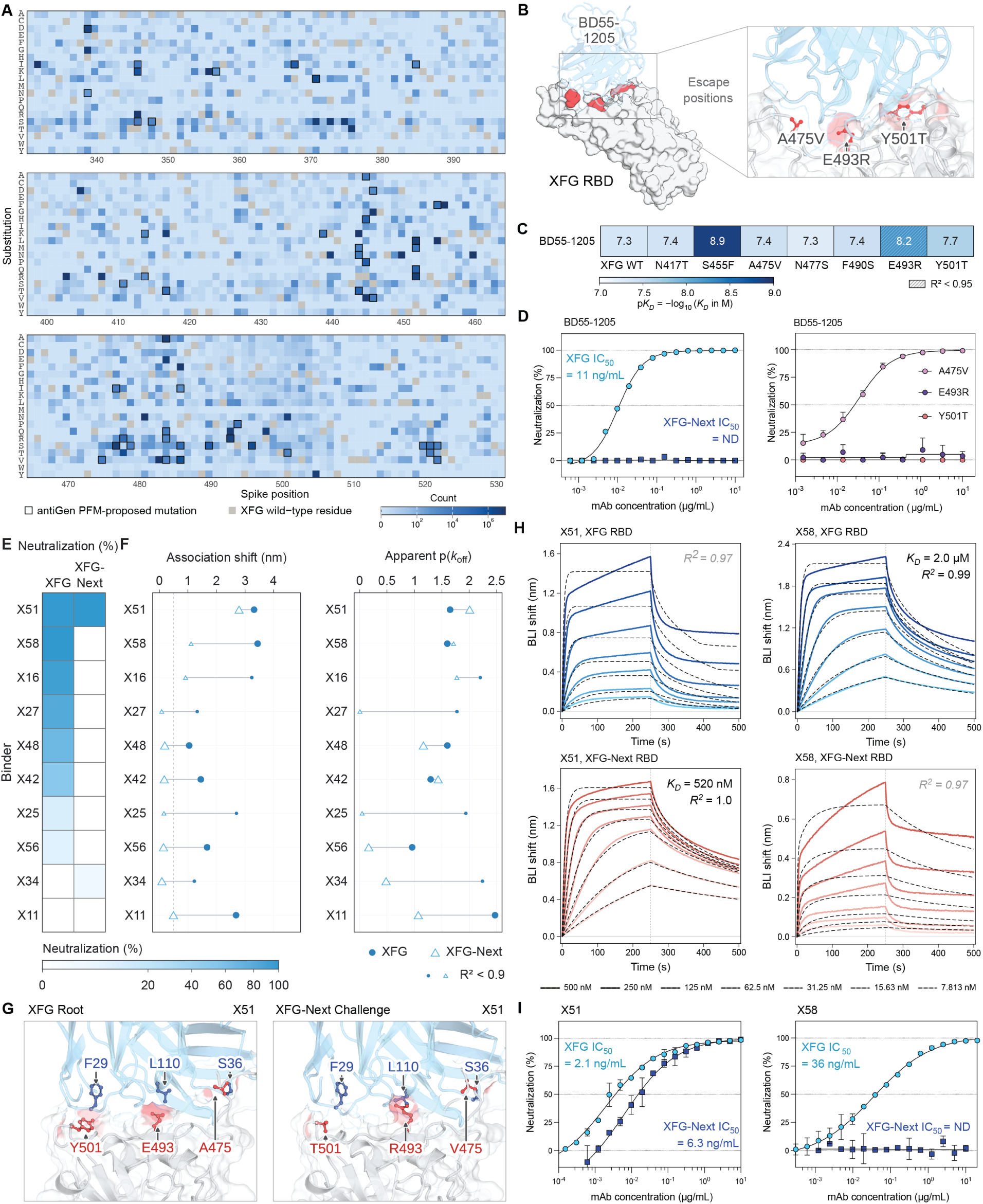
Antibody rescue against forecasted SARS-CoV-2 escape. **(A)** Amino-acid observation counts across the spike RBD. Outlined cells identify antiGen position frequency matrix (PFM)-proposed substitutions; gray cells mark the parental XFG residue at each position. **(B)** Structural view of BD55-1205 bound to the XFG RBD, highlighting the three positions substituted in the engineered escape variant XFG-Next (A475V, E493R, and Y501T). **(C)** Apparent *pK_D_* estimates for BD55-1205 binding to parental XFG and the indicated single-substitution RBD variants, where *pK_D_* = − log_10_ [*K_D_*/(M)]. Hatching indicates fits with *R*^2^ < 0.95. Corresponding sensorgrams are shown in **Figure S18**. **(D)** BD55-1205 neutralization of parental XFG and XFG-Next pseudoviruses (left) and XFG pseudoviruses carrying A475V, E493R, or Y501T individually (right). **(E)** Single-concentration neutralization results for the top ten of 60 screened antibody designs, ranked by mean percent neutralization across XFG and XFG-Next at 10 µg mL^−1^. Heatmap values show mean percent neutralization from two independent experiments, each performed in technical duplicate. **(F)** Single-concentration BLI screening of the ten designs shown in **(E)** against XFG and XFG-Next at 1,000 nM antibody. Plots show association responses and apparent dissociation rates expressed as *p*(*k*_off_) = − log_10_ [*k*_off_/(s^−1^)]; higher values indicate slower dissociation. Filled circles represent XFG; open triangles represent XFG-Next. Lines connect measurements for the same design; smaller symbols indicate kinetic fits with *R*^2^ < 0.9. Full sensorgrams and additional screening results are shown in **Figures S20, S21, and S10B**. **(G)** Structural models of X51 bound to XFG and XFG-Next RBDs. Selected antibody residues are labeled in blue; antigen residues at the three positions altered in XFG-Next are labeled in red. **(H)** BLI titrations of X51 and X58 against XFG and XFG-Next RBDs. Colored solid traces show reference-subtracted responses at the indicated antibody concentrations; black dashed traces show kinetic fits. Vertical dotted lines mark the start of dissociation at 250 s. Apparent *K_D_* estimates and corresponding *R*^2^ values are annotated for fits with *R*^2^ ≥ 0.99; otherwise, only *R*^2^ values are shown in gray italics. Kinetic fitting is described in **Methods**. **(I)** Neutralization of XFG and XFG-Next pseudoviruses by X51 and X58. For **(D,I)**, two independent experiments were performed, each with two technical replicates per concentration. Points show the mean of the two experiment-level means; error bars indicate SD (*n* = 2 independent experiments). Lines show neutralization curves fitted as described in **Methods**. Annotated IC_50_ values are the geometric means of the unrounded fitted estimates from two independent experiments, expressed in ng/mL. Experiment-specific fits for X51 and X58 are shown in **Figure S19**. ND indicates that an IC_50_ could not be determined within the tested concentration range.

We then assessed BD55-1205 recognition of the forecasted single-substitution variants in the XFG background. BD55-1205 retained measurable binding to all tested isolated RBD variants, although apparent affinities varied (**Figure 4C** and **Figure S18**). For substitutions outside the six DMS-defined escape-associated sites, we observed no apparent reduction in neutralization relative to parental XFG over the tested antibody concentration range (**Figure S17**).

Although BD55-1205 binding to the corresponding isolated RBDs remained measurable, variants carrying E493R or Y501T showed reduced susceptibility to BD55-1205 neutralization, whereas the A475V variant showed no measurable reduction despite its reported escape effect in the KP.3.1.1 DMS (**Figure 4D, right**). This difference is consistent with previously reported sequence-background dependence of antibody escape (Starr et al., 2022; Moulana et al., 2026).

We combined the three substitutions in the XFG background to generate the engineered challenge variant XFG-Next (A475V/E493R/Y501T). Its composition reflects the overlap between antiGen predictions and the published DMS landscape. BD55-1205 showed robust, concentration-dependent binding to parental XFG RBD but markedly reduced binding to XFG-Next RBD (**Figure S18**). Correspondingly, BD55-1205 neutralized parental XFG but did not show measurable neutralizing activity against XFG-Next within the tested antibody concentration range (**Figure 4D, left**), despite XFG-Next retaining measurable, albeit reduced, pseudoviral infectivity (**Figure S16**).

We next used Germinal Multi to redesign BD55-1205 to restore neutralization of the engineered escape variant XFG-Next while preserving activity against parental XFG. Unlike *de novo* design, rescue started from the parental antibody sequence, with a soft bias toward parental CDR residues. Structural templates of BD55-1205 bound to each target guided retention of its docking pose (**Methods**). This yielded 60 antibody candidates containing 11–20 CDR substitutions relative to BD55-1205 and predicted to bind both targets in the template docking pose.

Single-concentration neutralization screening identified X51 as a lead with activity against both XFG and XFG-Next (**Figure 4E and S10B**). We also screened the candidates for binding to both RBDs by BLI, applying the same hit criterion as above, i.e., a reference-subtracted response exceeding 0.5 nm at the end of the association phase against each RBD. Three designs met this criterion, with X51 and X58 producing the strongest association responses (**Figures S20, S21, and 4F**). Structural models illustrate the predicted interactions of X51 with XFG and XFG-Next (**Figure 4G**).

BLI titrations yielded comparable apparent *K_D_* estimates for X51 against XFG and XFG-Next, at 470 and 520 nM, respectively. X58 bound XFG-Next more strongly than XFG, with apparent *K_D_* estimates of 9.4 nM and 2.0 *μ*M, respectively (**Figure 4H**).

X51 neutralized both XFG (*IC*_50_ = 2.1 ng/mL) and XFG-Next (*IC*_50_ = 6.3 ng/mL), with approximately fivefold greater potency against XFG than the parental antibody BD55-1205 (*IC*_50_ = 11 ng/mL) (**Figure 4I and S19**). In contrast, X58 neutralized XFG (*IC*_50_ = 36 ng/mL) but showed no measurable neutralization of XFG-Next within the tested concentration range, despite binding both RBDs. These results underscore that RBD binding alone does not establish neutralizing activity.

We also assessed X51 against XFG variants carrying each constituent substitution of XFG-Next individually. X51 bound all three single-substitution RBD variants but neutralized only the A475V and E493R pseudovirus variants, with IC_50_ values of 47 and 69 ng/mL, respectively (**Figure S22A–C**, **S23**). However, X51 showed no detectable neutralization of Y501T within the tested concentration range (**Figure S23**). Together, these results demonstrate rescue of neutralization against the engineered challenge while preserving activity against parental XFG, but also show that activity against both targets does not ensure neutralization of every constituent single-substitution variant.

## 3. Discussion

In this study, we present a pipeline for *de novo* antibody design and rescue across viral variants and cancer-associated resistance variants, including an engineered escape variant informed by evolutionary forecasting. Across the design campaigns, selected antibodies achieved pseudoviral neutralization *IC*_50_ values in the low nanogram-per-milliliter range or apparent *K_D_* estimates in the single-digit nanomolar range. Successful candidates were identified after experimentally testing tens of designs, supporting the feasibility of extending computational antibody design from individual antigens to sets of antigenic variants. Computational design complements traditional antibody discovery by allowing specified antigen variants to be incorporated directly into the design objective. This provides a means of explicitly targeting breadth without requiring the desired recognition profile to be identified within an available donor repertoire.

An important question for future work is how the choice of design targets influences breadth, particularly against variants not explicitly included during optimization. In our forecast-informed campaign, target selection drew on evolutionary predictions and a DMS landscape from an earlier variant background, with subsequent experimental assessment in the contemporary background. This illustrates a practical approach to selecting targets when strain-matched experimental data are unavailable. More broadly, we applied Germinal Multi to evolutionary variants of the same protein; its applicability to more diverse target sets remains to be established. Although we focused on antibody breadth, a similar optimization strategy could support selective recognition of a desired subset of related antigens while limiting recognition of others.

The forecast-informed campaign also illustrates a potential application to pandemic preparedness and biodefense, where predicted amino acid changes can help assess antibody robustness and inform countermeasure development before such changes emerge in circulation. Here, evolutionary forecasting guided the design of an engineered variant, XFG-Next, that escaped an existing bnAb, and antibody redesign restored neutralization while retaining activity against the parental variant, XFG. Although the likelihood of XFG-Next emerging or spreading remains uncertain, the successful antibody rescue provides a proof of concept for integrating evolutionary forecasting with antibody design to support countermeasure development.

Recognition of multiple intended variants must also be distinguished from polyreactivity toward unrelated antigens. BLI screening detected no binding to the unrelated antigen PD-L1 under the tested conditions (**Figure S24**). Broader characterization against unrelated antigens or cellular material, together with assessment of biophysical properties, will be needed to evaluate the suitability of these designs for therapeutic development. Notably, our previous study describing the original Germinal method demonstrated both epitope targeting and low polyreactivity of *de novo* antibody designs (Mille-Fragoso et al., 2026).

Together, these results establish Germinal Multi as a framework for antibody co-optimization across defined sets of antigenic variants, supporting both *de novo* design and redesign of existing antibodies. We make our code and methods publicly available to support further evaluation and adoption of the approach, providing a foundation for proactive antibody development that incorporates robustness across specified antigenic variants as an explicit design goal.

## 4. Methods

### 4.1. Germinal1.1

Alongside the development of multi-target capabilities, we introduced several improvements to the original Germinal pipeline (Mille-Fragoso et al., 2026), resulting in an updated version termed *Germinal1.1*. These improvements were inspired by ProteinHunter (Cho et al., 2025), which demonstrated that iterative cycles of structure prediction and sequence redesign can quickly converge on binder designs with favorable *in silico* metrics. Notably, this work shows the initial structural proposal in this pipeline can be a weak design. Despite this, subsequent cycles of sequence redesign and structure prediction refine this initial proposal into designs with improved predicted structural properties.

The original Germinal pipeline, consists of three optimization stages—logit, softmax, and semi-greedy—that progressively anneal continuous logits into one-hot sequence representations. Inspired by ProteinHunter, we reasoned that the initial “logit” stage of hallucination in Germinal, despite not having a resolved sequence, can quickly generate the initial backbone proposal of the binder. In Germinal1.1, we adopt a similar procedure of iterative rounds of structural prediction of sequence design. Using only the logit stage yields a ∼2-fold speedup in proposal generation compared with standard Germinal1.

### 4.2. Multi-target co-optimization with Germinal Multi

Germinal Multi extends the Germinal1.1 framework to jointly optimize a shared antibody sequence against multiple targets (**Figure 1E**). Each target is supplied as a separate complex with its own template, sequence, and hotspot specification, while all complexes share a single set of designable antibody sequence logits. At each logit optimization step, AF2-Multimer is evaluated separately for each target, yielding target-specific losses and gradients with respect to the shared logits. Similar to concurrent work which co-optimized miniproteins against multiple variants (Brasas et al., 2026), gradients are combined either by summation or using PCGrad, which projects out conflicting gradient components before combining them. The recycling loop is generalized to multiple targets analogously: sequence redesign uses a multi-state ProteinMPNN formulation (with AbMPNN weights) in which both predicted complexes are supplied at once with symmetry constraints tying the binder positions across states, so a single sequence is decoded that is jointly consistent with both backbones, and the resulting sequence is folded separately against each target. A cycle is accepted only if all targets satisfy, hotspot-proximity, interface-CDR requirements, and the *minimum* ipSAE across targets exceeds the best value obtained previously. Rejected cycles revert to the best-scoring set of previously accepted backbones, so the minimum ipSAE of accepted design remains constant or increases monotonically.

### 4.3. Sequence and structure constrained multi-epitope co-optimization

For co-optimization of an antibody against XFG and the prospective XFG-Next receptor binding domain, instead of *de novo* antibody design, we opted to “rescue” binding of BD55-1205, an antibody which can recognize XFG but not the forecasted variant XFG-Next. To do this, two primary settings were altered to the pipeline. First, rather than initializing the antibody from a randomly sampled (Gumbel–softmax) sequence, we initialized directly from the parental BD55-1205 sequence and applied an additive logit bias toward the parental amino-acid identity at every CDR position (recover_bias = 5). This logit bias, termed (recover_bias = 5) is identical to the (framework_bias = 5) term used in Germinal, except applied to the CDRs. Conversely, in *de novo* co-optimization, these CDR biases are set to zero, leaving the loops free to explore. The bias acts as a soft restraint that makes the parental residue the default at each position while still allowing the optimizer to override it where the XFG-Next interface demands a substitution. Second, we provide a full-structural template of the parental complex against both targets to AF-M during design. Specifically, the BD55-1205–XFG and BD55-1205–XFG-Next structures were supplied as AlphaFold3 predicted input complexes, and inter-chain template contacts were retained (rm_template_ic = false) rather than removed as in *de novo* design. This means at every prediction iteration, AF2-M is given a template of the native BD55-1205 vs XFG dock. Together, the native sequence bias and native-dock template, provide the native sequence and structure as an explicit prior at both states.

### 4.4. Design filtering

Our design filters largely remain the same as our previous work (Mille-Fragoso et al., 2026), but are extended to evaluate on each target being optimized against. Additionally, we introduce several minor modifications to confidence filtering to enhance the stringency and reliability of calculations. First, we incorporate the ipSAE metric (Dunbrack, 2025), as it has been shown to be superior to ipTM at discriminating binders from non-binders (Overath et al., 2025). Second, given that AlphaFold3 (AF3) (Abramson et al., 2024) and PyRosetta (Chaudhury et al., 2010) are both non-deterministic functions, we run them for five seeds: for AF3 we take the confidence of the *best* diffusion sample (as is the typical behavior) but of the *worst* scoring seed, and for Rosetta we take the average of its metrics across seeds, as this has been shown to give more accurate results for protein property prediction (Wang et al., 2022). Finally, recognizing that antibodies can bind via a mixture of polar and hydrophobic interactions, or dominantly via either one, we combine our interface hydrogen-bond and interface hydrophobicity criteria into a single disjunctive filter rather than requiring both independently.

- AF3 pLDDT ≥ 0.85
- AF3 pTM ≥ 0.80
- AF3 ipTM ≥ 0.75
- AF3 PAE ≤ 9
- ipSAE ≥ 0.55
- clashes ≤ 0.2
- sc_rmsd ≤ 6 Å
- binder_near_hotspot == True
- cdr_interface_percentage ≥ 0.55
- interface_shape_complementarity ≥ 0.55
- surface_hydrophobicity ≤ 42%
- interface_hydrogen_bonds ≥ 8 **or** interface_hydrophobicity ∈ [40%, 60%]

### 4.5. Sequence and structure constrained filtering

For XFG-Next binders derived from BD55-1205, we applied additional filters to favor retention of the parental binding mode. Candidates were required to contain no more than 20 amino-acid substitutions relative to parental BD55-1205 and to satisfy the structural and confidence thresholds used in the final filtering stage. We computed the C*α* RMSD of each predicted complex—with XFG or XFG-Next—relative to the parental BD55-1205:XFG template complex and retained designs only if the maximum of the two RMSD values were below 5 Å. The resulting pool was ranked by ipSAE, and the top 60 designs were selected for downstream experimental characterization.

### 4.6. Generation of ESM-IF-, FAMPNN- and MSA-guided AZD3152 benchmarking variants

To benchmark model-guided rescue within the local mutational neighborhood of AZD3152, we generated single-amino-acid substitution panels using ESM-IF, FAMPNN and an AbYsis-derived multiple-sequence-alignment (MSA) baseline (Shanker et al., 2024; Hsu et al., 2022; Widatalla et al., 2025). Structure-conditioned proposals were generated independently in the BA.2 root and KP.3 challenge contexts, both built from the experimentally determined AZD3152–BA.2 complex (PDB 8SUO) (Cai et al., 2024). The root context used the deposited complex directly; the challenge context was constructed by substituting the KP.3 receptor-binding-domain sequence onto the same backbone at the fifteen positions that differ from BA.2.

#### ESM-IF-guided mutation generation

ESM-IF1 was used to score single-amino-acid substitutions conditional on the antibody–antigen complex structure, following the whole-complex inverse-folding procedure described previously (Shanker et al., 2024; Hsu et al., 2022). For a complex with backbone coordinates **Y** and sequence **x**, ESM-IF estimates the conditional sequence likelihood of a given variant sequence *p*(**x** | **Y**). All possible non-parental amino acids were introduced individually at each AZD3152 VH and VL position and the resulting complex re-scored; the antigen sequence was held fixed within each context. Scoring used the released ESM-IF1 weights (esm_if1_gvp4_t16_142M_UR50), conditioned jointly on all three chains of the complex.

The BA.2- and KP.3-bound complexes were scored independently, producing root-conditioned and challenge-conditioned rankings. Candidate substitutions were ranked by the mutant-minus-parental difference in whole-chain conditional log-likelihood, log *p*(**x**_mut_ | **Y**) − log *p*(**x**_wt_ | **Y**), evaluated over the entire mutated antibody chain, and the five highest-ranked substitutions at unique heavy-chain positions and the five highest-ranked substitutions at unique light-chain positions were selected from each antigen context. Thus, ESM-IF contributed four proposal sets: heavy- and light-chain substitutions conditioned on BA.2, and heavy- and light-chain substitutions conditioned on KP.3.

#### FAMPNN-guided mutation generation

FAMPNN was used as a complementary full-atom inverse-folding model (Widatalla et al., 2025). In contrast to ESM-IF, which conditions primarily on protein-backbone geometry, FAMPNN incorporates the local all-atom environment, including side-chain coordinates, when evaluating sequence compatibility.

Each substitution was scored by masking the residue identity and all side-chain coordinates at the candidate position and taking the difference in the model’s log-probability at that position, log *p*(*a*_mut_) −log *p*(*a*_wt_), conditioned on the surrounding all-atom environment. Because the challenge context changes the antigen sequence on a fixed backbone, antigen side chains were re-predicted by FAMPNN before scoring: the receptor-binding domain was repacked with the model’s side-chain diffusion module while the antibody side chains were retained as deposited. Fifty packed structures were generated, the five ranking best on combined steric-clash and predicted side-chain-error percentile were retained, and per-substitution scores were averaged across them.

All possible non-parental single substitutions were evaluated separately in the root and challenge complexes. Candidate substitutions were ranked independently for VH and VL, and the five highest-ranked substitutions at unique positions in each chain were selected from each antigen context. FAMPNN therefore contributed an additional four proposal sets: BA.2-conditioned heavy- and light-chain substitutions and KP.3-conditioned heavy- and light-chain substitutions.

#### AbYsis MSA-guided mutation generation

An antigen-independent evolutionary baseline was generated using position-specific amino-acid frequencies from AbYsis webtool version 3.4.1 (http://www.abysis.org/abysis/index.html). The AZD3152 VH and VL sequences were aligned using the AbYsis Annotate tool under its default settings against a database of human antibody sequences. For each aligned variable-domain position, AbYsis reports the frequency of each amino acid observed among naturally occurring antibody sequences.

Each possible non-parental substitution was assigned the frequency of the mutant residue at the corresponding aligned position. Substitutions were ranked independently within VH and VL, and the five highest-ranked heavy-chain and five highest-ranked light-chain substitutions at unique positions were selected. Because this baseline did not use an antigen structure or sequence, a single MSA proposal set was applied to both the BA.2 and KP.3 binding objectives.

### 4.7. Position frequency matrix (PFM) forecasting of SARS-CoV-2 XFG RBD evolution

We forecasted the 50 most likely single amino acid substitutions relative to the XFG wild-type RBD using a position frequency matrix (PFM)-based approach (Specht et al., 2026). Using all public SARS-CoV-2 sequences collected over the first five years of the pandemic, retrieved through the outbreak.info API (Gangavarapu et al., 2023), we counted the number of observations of amino acid *i* at position *j*, for *i* ranging over all RBD positions and *j* over all amino acids. We then zeroed out the count for any pair (*i*, *j*) for which the XFG wild-type amino acid was equal to *j*. At the time of analysis, we extracted the 50 amino acid substitutions with the highest remaining counts and deposited the results in Zenodo 10.5281/zenodo.17877515. Ranking table and PFM scores can be found in **Data S1**. Previous analysis (Specht et al., 2026) has demonstrated this counting method to be a very strong predictor of future mutations.

Predicted substitutions were further restricted to those i) within the BD55-1205 binding interface, defined by superimposing the predicted XFG–BD55-1205 complex onto the experimentally determined XBB.1.5–BD55-1205 complex (Jian et al., 2025), and ii) at sites associated with high BD55-1205 neutralization escape and retained ACE2 binding in published pseudovirus-based deep mutational scanning measurements obtained in a KP.3.1.1 spike background (Dadonaite et al., 2025). Three candidate substitutions–A475V, E493R, and Y501T–were evaluated for their effects on BD55-1205 binding by BLI and on spike-mediated cell entry using XFG pseudovirus infectivity assays. The three substitutions were combined to generate the engineered challenge antigen termed XFG-Next.

For supporting surveillance context, daily counts of sequences carrying A475V obtained from the LAPIS API (Chen et al., 2023) and the corresponding 30-day centered rolling mean are shown in **Figure S15**.

### 4.8. Plasmids and cloning

Designed antibody and recombinant antigen coding sequences were codon-optimized for human expression and synthesized by Twist Bioscience. SARS-CoV-2 RBD constructs for recombinant protein production were based on spike protein sequences with the following GenBank protein accession numbers: BA.2 (XXK88105.1), BA.2.86 (XAT77441.1), JN.1 (XPR37317.1), JN.1.11.1 (XPZ25006.1), KP.3 (XRH93350.1), PG.3.1 (XQO33767.1), and XFG (YCJ32983.1).

Expression constructs were assembled by PCR using NEBNext High-Fidelity polymerase (New England Biolabs, M0544) followed by Gibson assembly (New England Biolabs, E2621S) or In-Fusion assembly using In-Fusion Snap Assembly Master Mix (Takara Bio, 638944). Synthetic fragments encoding recombinant antigens were inserted into pCMV-derived mammalian expression vectors containing an N-terminal signal peptide, a C-terminal AviTag (GLN-DIFEAQKIEWHE), and a His_6_-tag. RBD constructs used the SARS-CoV-2 spike signal peptide (MFVFLVLLPLVSSQ), whereas EGFR-Fc constructs used the native signal peptide (MRPSGTAGAALLALLAALCPASRA). Fragments encoding antibody variable heavy-chain (VH) regions were cloned into human IgG1 heavy-chain expression plasmids. Fragments encoding antibody variable light-chain (VL) regions were cloned into either a human kappa light-chain expression plasmid (for cetuximab, 13D4, BD55-1205, and designed binders using the BD55-1205 or trastuzumab frameworks) or a human lambda light-chain expression plasmid (for AZD3152). Both heavy- and light-chain expression vectors contained an N-terminal signal peptide derived from the murine immunoglobulin heavy chain (MGWSCIILFLVATATGVHS) to direct antibody secretion. Sequences and expression vectors of the constructs described in this study are provided in **Data S2, S3**. Individual mutations guided by MSA, ESM-IF, or FAMPNN, along with combinations of these mutations, were introduced into the AZD3152 variable heavy- or light-chain domain sequences by site-directed mutagenesis using primers listed in (**Data S3**).

For SARS-CoV-2 pseudovirus production, the Wuhan-Hu-1 HDM-SARS2-Spike-Δ21 expression plasmid (Addgene 155130) was used as the parental backbone. Nucleotides 1364–4745 of the Wuhan-Hu-1 spike sequence was replaced with the corresponding BA.2, KP.3 or XFG spike sequence by PCR-based cloning. Individual substitutions generating the XFG-Next spike were introduced by site-directed mutagenesis (**Data S2**). The packaging plasmids were pHAGE-Luc2-IRES-ZsGreen (BEI Resources NR-52516), HDM-Hgpm2 (NR-52517), HDM-Tat1b (NR-52518) and pRC-CMV-Rev1b (NR-52519) (**Data S2**). Expression plasmids for spike proteins, antibodies, and antigens were propagated in Stellar competent cells (Takara Bio, 636766), and all final constructs were validated by whole-plasmid Nanopore sequencing.

### 4.9. Cell culture

Expi293F cells (Thermo Fisher Scientific, A14635) were maintained in a 2:1 mixture of FreeStyle 293 Expression Medium (Thermo Fisher Scientific, 12338018) and Expi293 Expression Medium (Thermo Fisher Scientific, A1435102) at 37 ^◦^C, 8% CO_2_ and 125 rpm. HEK293T pseudovirus producer cells were obtained from John Long’s laboratory at Stanford University and HEK293T/ACE2/TMPRSS2 target cells were obtained from Peter S. Kim’s laboratory at Stanford University. Both cell lines were maintained at 37 ^◦^C in a humidified incubator with 5% CO_2_ in complete D10 medium, consisting of Dulbecco’s Modified Eagle Medium (DMEM; Corning) supplemented with 10% fetal bovine serum (Thermo Fisher Scientific), 2 mM L-glutamine (Gibco), and 1% penicillin–streptomycin (Gibco).

### 4.10. Expi293 protein expression

Control antibodies were expressed in Expi293F cells (Thermo Fisher; #A14635). AviTag-fused antigens were expressed in Expi293F cells in the presence of BirA to enable their*in vitro* biotinylation for BLI experiments. The day before transfection, Expi293F cells were diluted to 2 × 10^6^ cells mL^−1^ and cultured for 20–24 h. Cells were then adjusted to 3 × 10^6^ cells mL^−1^ for transfection in final culture volumes of 30–250 mL. Transfection mixtures contained 0.5 µg plasmid DNA, 1.3 µL FectoPRO transfection reagent (Polyplus, 101000007) and 100 µL expression medium per millilitre of culture. For IgG expression, heavy- and light-chain plasmid DNA were combined at a 1:2 ratio. Transfection mixtures were incubated for 10 min before addition to cells. Cultures were immediately supplemented with glucose to 0.45% (w/v) and valproic acid to 2 mM, then cultured for 5 d. Cultures expressing AviTag-fused antigens were additionally supplemented with biotin to 5 mM (Invitrogen; B20656).

For higher-throughput expression of MSA-, ESM-IF-, or FAMPNN-informed antibody mutants, Expi293F cells were prepared as described above and seeded at 3 × 10^6^ cells mL^−1^ in round-bottom 96-well deep-well plates at 1 mL per well on the day of transfection. Heavy- and light-chain plasmid DNA were combined at a 1:3 ratio. Following transfection, plates were incubated for 5 d at 37 ^◦^C, 8% CO_2_ and 1000 rpm with a 3 mm (diameter) orbital throw.

Culture supernatants were clarified by centrifugation at 4000 ×*g* for 15 min at 4 °C and filtered through a 0.22 *μ*m syringe filter (Thermo Fisher; 723-2520). For antibodies expressed in 96-well plates, cells were pelleted by centrifugation and supernatants were carefully transferred to fresh plate without disturbing the cell pellets. Filtered supernatants were supplemented with 10× phosphate buffered saline (PBS; 1.37 M NaCl, 27 mM KCl, 100 mM Na2HPO4, 18 mM KH2PO4, pH 7.4) and 1.63 M NaCl, then either purified immediately or stored at −20 ^◦^C for less than 1 month.

### 4.11. Antibody expression and purification using the TurboCHO

Designed antibodies were expressed as full-length IgG by GenScript using TurboCHO™ Express Platform. The heavy- and light-chain variable regions from each scFv design were incorporated into an a human IgG1 backbone, retaining the corresponding *κ* or *λ* light-chain type. Heavy- and light-chain sequences were codon-optimized, synthesized, and cloned into the platform’s proprietary expression vectors, then transiently transfected into suspension-adapted CHO cells grown in chemically defined, serum-free medium. The specific host cell line, transfection method, and culture conditions were proprietary and were not disclosed by the vendor. Secreted antibodies were purified from culture supernatants by Protein A affinity chromatography without additional purification tags. Designs were assessed by SDS-PAGE (**Figure S28**) and size-exclusion high-performance liquid chromatography (SEC-HPLC).

### 4.12. Antigen purification by immobilized metal affinity chromatography and size-exclusion chromatography

His-tagged recombinant antigens were purified using HisPur Cobalt resin (Thermo Fisher Scientific, 89965). Resin was washed twice in binding buffer (300 mM NaCl, 2.7 mM KCl, 10 mM Na_2_HPO_4_ and 1.8 mM KH_2_PO_4_, pH 7.4) supplemented with 10 mM imidazole, then added to clarified culture supernatant and incubated for 1 h at 4 ^◦^C with end-over-end rotation. The mixtures were transferred to gravity-flow columns and washed sequentially with binding buffer and binding buffer supplemented with 30 mM imidazole. Proteins were eluted in binding buffer containing 200 mM imidazole.

Elutes were immediately concentrated and buffer-exchanged into PBS, pH 7.4 using centrifugal concentrators with 5- or 10-kDa molecular-weight-cutoffs. Buffer exchange was performed over four centrifugation cycles at 4,000-5,000 ×*g* for 10 min at 4 ^◦^C, with dilution in PBS between cycles.

All antigen preparations were further purified by ÄKTA pure fast-protein-liquid-chromatography system (FPLC; Cytiva) equipped with Superdex 200 Increase 10/300 GL column pre-equilibrated with PBS, pH 7.4. A sample loading volume of 0.5 mL and flow rate of up to 1 mL/min was used. Peak fractions were assessed by SDS-PAGE (**Figure S28**), and fractions containing the target species were pooled and concentrated. Protein concentrations were determined by absorbance at 280 nm using sequence-specific extinction coefficients calculated with ExPASy ProtParam. Purified proteins were flash-frozen in liquid nitrogen and stored at −80 ^◦^C. Antigen size-exclusion chromatograms are shown in **Figure S25** for the EGFR-Fc panel, **Figure S26** for the BA.2-derived RBD panel, and **Figure S27** for XFG and XFG-Next.

### 4.13. Antibody purification by Protein A affinity chromatography

IgG antibodies expressed in volumes greater than 30 mL were purified by Protein A affinity chromatography using a 5-mL HiTrap MabSelect PrismA column (Cytiva; 17549853) connected to an ÄKTA pure FPLC system (Cytiva). The system and column were equilibrated with PBS, pH 7.4. Clarified, filtered Expi293F culture supernatants were loaded from a 50-mL Superloop at a flow rate of 2.5 mL min^−1^. The column was washed with at least five column volumes of PBS pH 7.4, and bound antibodies were eluted with 20 column volumes of 0.1 M sodium citrate, pH 3.0. Eluted fractions were collected directly into tubes containing 1 M Tris-HCl pH 9.0, for immediate neutralization. Peak fractions were assessed by SDS-PAGE (**Figure S28**), and antibody-containing fractions were pooled and concentrated using centrifugal concentrators with 10- or 30-kDa molecular-weight cutoffs. Purified antibodies were either used directly in downstream experiments or flash-frozen and stored at −80 ^◦^C.

Between purification runs for different antibody samples, the column was equilibrated with seven column volumes of PBS, pH 7.4, washed with two column volumes of 2.5 M NaCl, and re-equilibrated in PBS, pH 7.4.

### 4.14. Antibody purification using Protein G magnetic beads

IgG antibodies expressed in 96-well deep-well plates were purified using Magne Protein G magnetic beads (Promega; V1605). Protein G magnetic beads were washed twice in PBS, pH 7.4, and resuspended in PBS as a 20% slurry. A 50 *μ*L washed beads were added to each well containing clarified culture supernatant and incubated for 30–60 min at room temperature on a plate shaker at 900-1,000 rpm. Plates were placed on a magnetic stand for approximately 3 min, and unbound supernatants were removed without disturbing the pelleted beads. Beads were washed once with 500 µL PBS, pH 7.4, followed by two washes with 200 µL PBS, pH 7.4. For each wash, beads were fully resuspended before magnetic separation and removal of the wash buffer.

Bound antibodies were eluted by adding 50 µL of 100 mM glycine, pH 2.7, per well. Beads were resuspended by pipetting for approximately 30 s and incubated for 5 min at room temperature on a plate shaker at 900-1,000 rpm. Following magnetic separation using a magnetic plate, elutes were transferred to a fresh plate and immediately neutralized with 10 µL of 1 M Tris-HCl pH 9.0. Elution was repeated twice more, for a total of three elutions. Purified antibodies were used directly in downstream assays or buffer-exchanged into PBS as required.

CoV2-26, an antibody targeting the S2 subunit of the SARS-CoV-2 spike protein was a kind gift from the laboratory of Peter S. Kim and served as a negative control for RBD binding (Utz et al., 2025).

### 4.15. SDS-polyacrylamide gel electrophoresis (SDS-PAGE)

Protein samples were mixed with 4× Laemmli loading buffer to a final concentration of 1× (62.5 mM Tris–HCl, pH 6.8, 2% (w/v) SDS, 10% (v/v) glycerol and 0.002% (w/v) bromophenol blue) and denatured by boiling at 99 ^◦^C for 3 min. All antibody preparations, wild-type EGFR-Fc and EGFR-Fc mutants were analyzed under reducing and non-reducing conditions, with or without a final concentration of 2.5% (v/v) *β*-mercaptoethanol, respectively, to assess disulfide-linked assembly. Samples were resolved at 200 V on 4–12% SurePAGE Bis-Tris precast gels (GenScript, M00654) in an XCell SureLock Mini-Cell apparatus (Thermo Fisher Scientific, EI0001) using 1× Tris–MOPS–SDS running buffer (GenScript, M00138). PageRuler Plus (Thermo Fisher Scientific, 26619) was used as the molecular-weight standard. Gels were stained with InstantBlue Coomassie (Abcam, ab119211) with gentle agitation for up to 1 h, destained in Milli-Q water for at least 1 h and imaged using a GelDoc Go system (Bio-Rad). Images were analyzed in ImageLab version 6.1.0.

### 4.16. Production of SARS-CoV-2 pseudovirus

Replication-incompetent SARS-CoV-2 spike-pseudotyped lentiviral particles were generated in HEK293T cells using a five-plasmid system. HEK293T producer cells were seeded at 6 × 10^6^ cells in 10 mL per 10-cm dish one day before transfection. Per dish, plasmids were mixed in a final volume of 1 mL DMEM containing 10 µg pHAGE-Luc2-IRES-ZsGreen, 3.4 µg spike-expression plasmid, 2.2 µg HDM-Hgpm2, 2.2 µg HDM-Tat1b, and 2.2 µg pRC-CMV-Rev1b (Crawford et al., 2020). Then, 30 µL of BioT transfection reagent (BioLand, B01-01) was added to the plasmid mixture, and the mixture was incubated for 10 min at room temperature. Subsequently, 9 mL D10 medium was added to the mixture to bring the final volume to 10 mL, and the resulting transfection mixture was added to the cells (Powell et al., 2026).

After 24 h, the medium was replaced with 10 mL of fresh D10. Pseudovirus-containing supernatant was harvested 72 h after transfection, centrifuged at 300 ×*g* for 5 min and filtered through a 0.45-µm PES filter. Stocks were aliquoted and stored at −80 ^◦^C. When concentrated stocks were required, filtered supernatants were mixed with Lenti-X Concentrator (Takara Bio, 631231) at a 3:1 supernatant-to-reagent ratio, incubated overnight at 4 ^◦^C, and centrifuged at 1,500 ×*g* for 45 min. Pellets were resuspended in D10 at one-tenth of the original supernatant volume.

### 4.17. Pseudovirus entry and neutralization assays

In order to evaluate the effect of selected antiGen-PFM-forecasted mutations on host cell entry, we performed titrated infectivity assays (Figure S16). HEK293T/ACE2/TMPRSS2 target cells were seeded at 5 × 10^3^ cells per well. Pseudovirus preparations were serially diluted in D10 and added to cells at a total volume of 100 µL per well. Cells were incubated for 48 h at 37 ^◦^C before the addition of 100 µl BriteLite reagent prepared as a 1:1 mixture with 1x DPBS. Luminescence was measured in relative luminescence units (RLU) using a Tecan Spark multimode plate reader. We note that pseudovirus preparations were not normalized by particle number, RLU per µL of values were interpreted only as evidence that a substitution retained detectable spike-mediated entry, rather than a quantitative comparison of intrinsic infectivity.

For antibody neutralization, pseudovirus was incubated with antibody at 37 ^◦^C for 30 min before addition to target cells. To identify lead designs against BA.2–KP.3 and XFG–XFG-Next, an initial screen was performed at a final antibody concentration of 10 µg mL^−1^. BD55-1205, the parental antibody used in the XFG/XFG-Next design campaign, was used as a positive control for BA.2, KP.3, and XFG (Mellis et al., 2025; Jian et al., 2025). AZD3152, which neutralizes BA.2 but not KP.3 (Planas et al., 2024), was also included. The anti-H5 antibody 13D4 was used as the isotype control (Zheng et al., 2011).

Single-point screens were used only to identify lead candidates for further characterization and were not used to infer half-maximal inhibitory concentrations. Candidates that demonstrated at least 50% neutralization against both their root and challenge antigen target at 10 µg mL^−1^, relative to background-subtracted signals from virus-only control cells, were evaluated using the full antibody titrations to determine half-maximal inhibitory concentrations (IC_50_). Percent inhibition was calculated as [1 - (mAb-treated - cells-only)/(virus-only - cells-only)] × 100. IC_50_ values were determined by nonlinear regression using a four-parameter logistic (4PL) dose-response model with a variable slope. Values were reported as not determined (ND) when inhibition did not reach 50% within the tested concentration range (up to 50 µg mL^−1^). Normalized inhibition values were analyzed without truncation at 0% or 100%. Two independent experiments were performed, each with two technical replicates. Individual experiment IC_50_ values and their geometric mean were reported to two significant figures. Geometric means were calculated using the original, unrounded fitted IC_50_ values.

### 4.18. Avidity characterization by biolayer interferometry (BLI)

BLI experiments were performed on an Octet RED96 instrument at 30 ^◦^C in PBS, pH 7.4, supplemented with 0.2% BSA and 0.05% Tween-20 (assay buffer). Streptavidin Biosensors (Sartorius, 18-5019) were pre-incubated in assay buffer for 10 min and equilibrated to baseline before loading with biotinylated target antigens. For kinetic measurements, targets were loaded from 100 nM solutions for 300 s or until a loading response of 1 nm was reached. Loaded sensors were equilibrated to baseline in assay buffer before transfer to antibody solutions for association (200-250 s), followed by transfer to assay buffer for dissociation (200-250 s). A target-loaded sensor incubated in assay buffer without antibody served as the reference channel. Reference signals were subtracted from all sample channels before kinetic fitting.

For candidate screening, designs were initially evaluated for binding to their respective root targets (BA.2 or XFG) and challenge targets (KP.3 or XFG-Next), except for EGFR-Fc designs, which were tested only against wild-type EGFR-Fc. Binders with an association response greater than 0.5-nm against either target were subsequently titrated against their cognate antigens to obtain apparent avidity estimates.

For avidity characterization, four-to-seven antibody concentrations were prepared by two-fold serial dilution from the highest concentration indicated on each plot. Reference-subtracted sensorgrams were analyzed using Octet Data Analysis software (version 9.0.0.15) and globally fitted to a 1:2 bivalent model for IgG. Apparent *K_D_* estimates and the corresponding *R*^2^ values are shown on the respective sensorgram plots. Apparent *K_D_* estimates from fits with *R*^2^ < 0.99 are shown in gray italics.

### 4.19. Ethics declaration and institutional review

Work with SARS-CoV-2 spike-pseudotyped lentiviral particles was approved by the Stanford University Institutional Biosafety Committee/Administrative Panel on Biosafety under APB protocol 6007-BH0126. The protocol covers production and use of lentiviral particles pseudotyped with SARS-CoV-2 spike proteins from variant spike proteins, including emerging variants of concern, and spike constructs containing point mutations within the receptor-binding domain. All pseudotyped lentiviral particles were replication-incompetent, used for single-cycle entry infectivity assays, and handled under approved institutional biosafety procedures at BSL-2 containment.

### 4.20. Statement on AI use

Claude Opus 4.6–5 and GPT-5.5/6 Astra were used to assist in development of the codebase and sampling runs. This included launching experiments to compare varying design parameters, configuring jobs for sampling designs against the targets, parsing and preparing input structure and MSA files, preparing plotting code, and conducting checks for errors in the codebase. All authors take full responsibility for the entire content of the manuscript, including any portions developed with the assistance of AI tools, and have reviewed and verified all text, code, figures, and results.

## Supporting information

Data S1_XFG_RBD_predictions_PFM_counts

Data S2_Antigens_Pseudovirus_Information

Data S3_Antibodies

## 5. Data availability

A ranked list of the 50 most likely single amino acid substitutions relative to the XFG wild-type RBD, predicted using a position frequency matrix (PFM)-based approach, along with their corresponding PFM scores, is provided in **Data S1**. Sequences of antigen variants generated for BLI, viral variants used for pseudovirus production, and primers used for cloning, together with information on template vectors used for pseudovirus preparation, are provided in **Data S2**. Amino acid sequences of the antibody heavy- and light-chain variable regions used in this study, along with primer sequences used to generate the AZD3152 baseline mutants, are provided in **Data S3**. All three data files are available at https://doi.org/10.5281/zenodo.24030408.

## 6. Code availability

We make our code open to the public, with code for running Germinal Multi, including the configuration files specifying the parameters used for the three campaigns presented in this work, available at https://github.com/twidatalla/germinal-multi.

## 7. Acknowledgements

We thank members of the Laboratory of Evolutionary Design for helpful discussions and assistance with manuscript preparation. We thank members of the Peter S. Kim lab and John Pluvinage lab for contributing experimental reagents related to the pseudoviral infectivity assays and Biolayer Interferometry (BLI) measurements. S.P. acknowledges funding support from the Stanford Graduate Fellowship and Sarafan ChEM-H Chemistry/Biology Interface Program. T.W. acknowledges support by the NSF Graduate Research Fellowship and the Stanford Graduate Fellowship. L.S.M.-F. acknowledges support from the Stanford Interdisciplinary Graduate Fellowship and Stanford Bio-X. I.S. acknowledges funding support from the Fannie and John Hertz Foundation and Knight-Hennessy Scholars. B.L.H. acknowledges funding support from the Gates Foundation, the Silicon Valley Community Foundation, the NIH Director’s New Innovator Award, the Advanced Research Projects Agency for Health (ARPA-H), Arc Institute, Schmidt Sciences AI2050, Stanford Center for Digital Health, and Stanford Human-Centered Artificial Intelligence (HAI) Hoffman-Yee Research Grants.

## 8. Author contributions

T.W., C.L.D., L.S.M.-F., and B.L.H. conceived the project. C.L.D. and B.L.H. supervised the project. T.W., C.L.D., and L.S.M.-F. developed Germinal1.1 and Germinal Multi. T.W., C.L.D., and L.S.M.-F. conducted Germinal inference for antibody design. S.P., T.W., C.L.D., and L.S.M.-F. performed protein expression and purification. S.P., T.W., and C.L.D. performed antibody binding experiments. S.P. performed the SARS-CoV-2 neutralization experiments. I.S. conducted evolutionary forecasting with antiGen. S.P., T.W., C.L.D., L.S.M.-F., and B.L.H. wrote the initial draft of the manuscript. All authors contributed to the final draft of the manuscript.

## A. Supplementary Figures

**Figure S1.**
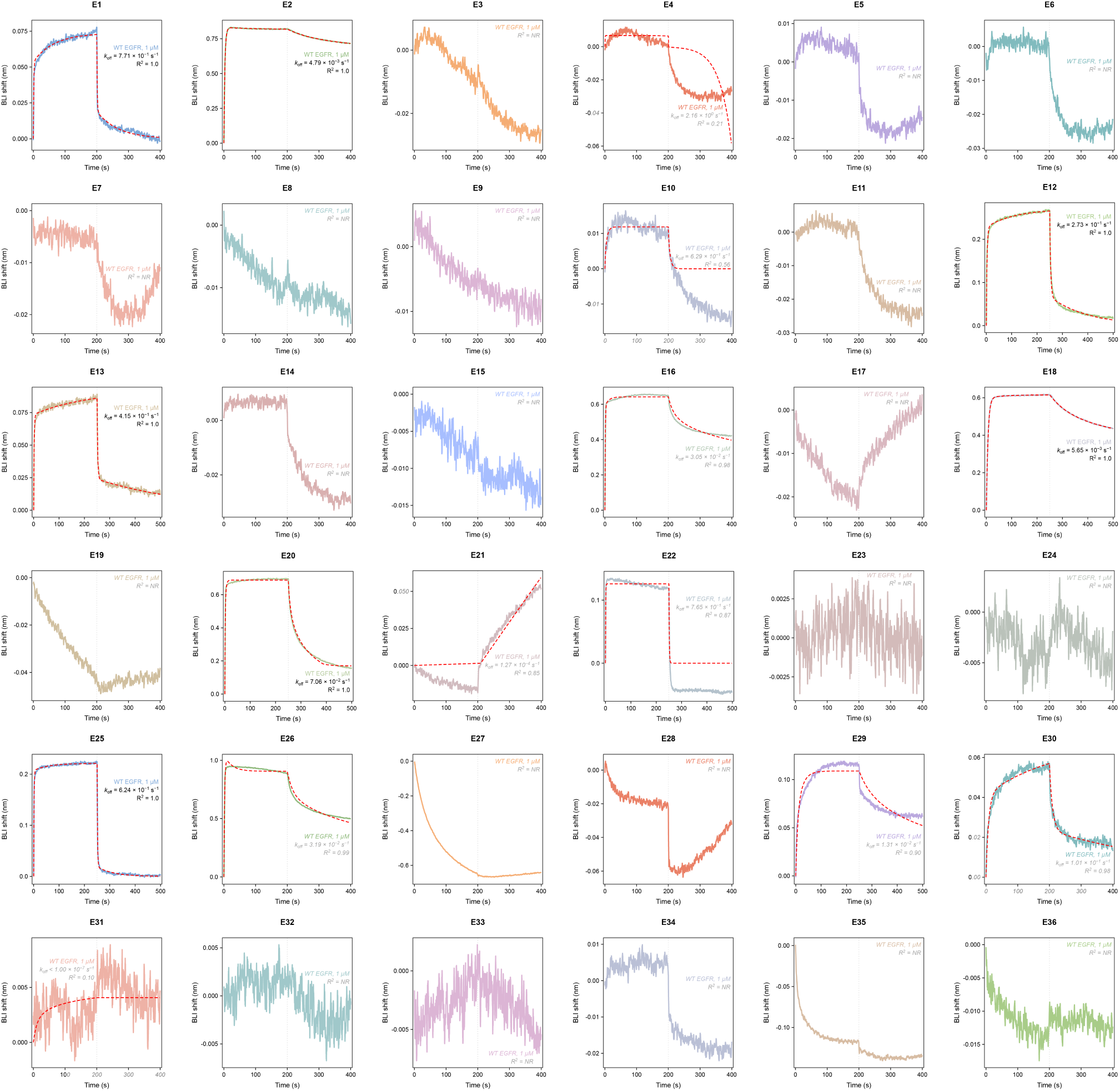
Single-concentration binding screen of EGFR designs E1–E36. Biolayer interferometry (BLI) sensorgrams for designs E1–E36 tested at 1,000 nM antibody against immobilized wild-type EGFR-Fc. Colored solid traces show reference-subtracted responses, and red dashed traces show kinetic fits where available. Dissociation rate constants (*k_off_*) were estimated from single-concentration screening measurements; corresponding *R*^2^ values are shown. Estimates from fits with *R*^2^ < 0.99 are shown in gray italics. Blank entries indicate that *k_off_* could not be determined; NR, not reported.

**Figure S2.**
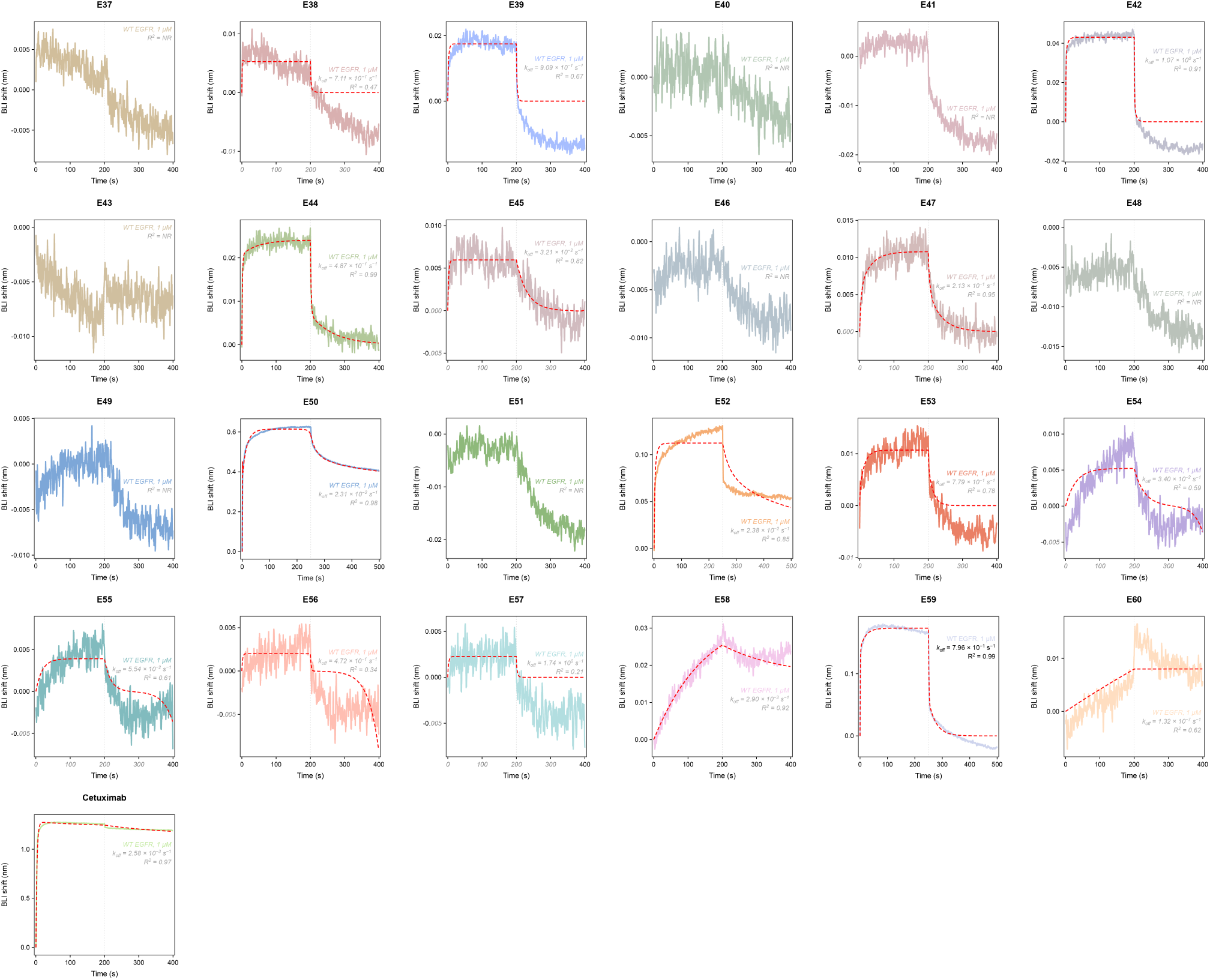
Single-concentration binding screen of EGFR designs E37–E60 and cetuximab. BLI sensorgrams for designs E37–E60 and the cetuximab control tested at 1,000 nM antibody against wild-type EGFR-Fc. Colored solid traces show reference-subtracted responses, and red dashed traces show kinetic fits where available. Dissociation rate constants (*k_off_*) were estimated from single-concentration screening measurements; corresponding R2 values are shown. Estimates from fits with *R*^2^ < 0.99 are shown in gray italics. Blank entries indicate that *k_off_* could not be determined; NR, not reported.

**Figure S3.**
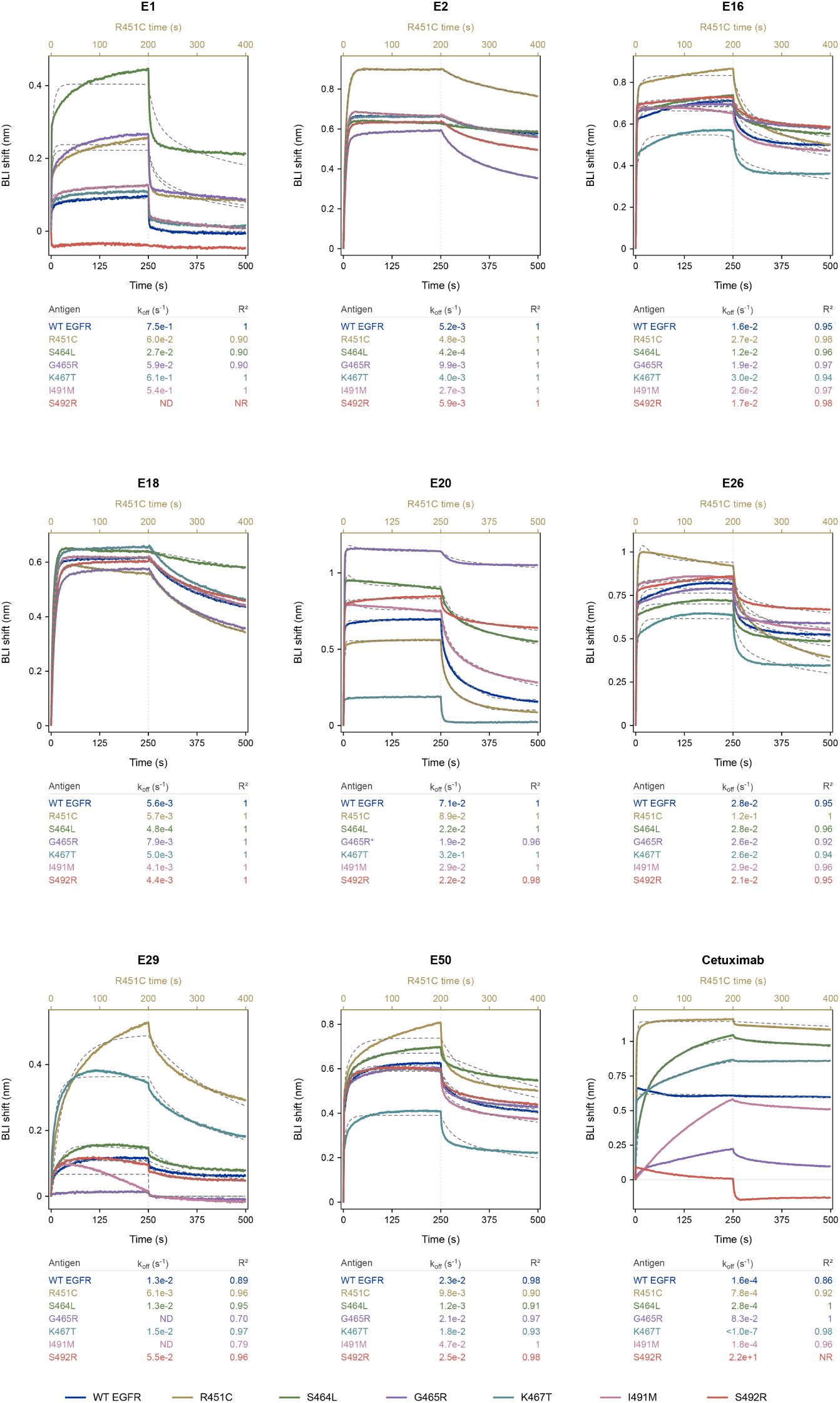
Binding profiles of selected EGFR designs against individual resistance variants. Single-concentration BLI measurements of E1, E2, E16, E18, E20, E26, E29, E50, and cetuximab against wild-type EGFR and the individual R451C, S464L, G465R, K467T, I491M, and S492R variants. Antibodies were tested at 1,000 nM. Colors identify antigens; gray dashed traces show kinetic fits where available. Tables report dissociation rate constant (*k*_off_) estimates and corresponding *R*^2^ values. R451C traces use the upper time axis to align association and dissociation phases with the other measurements; all other traces use the lower time axis. Estimates from fits with *R*^2^ < 0.99 are shown in gray italics. ND, not determined; NR, not reported.

**Figure S4.**
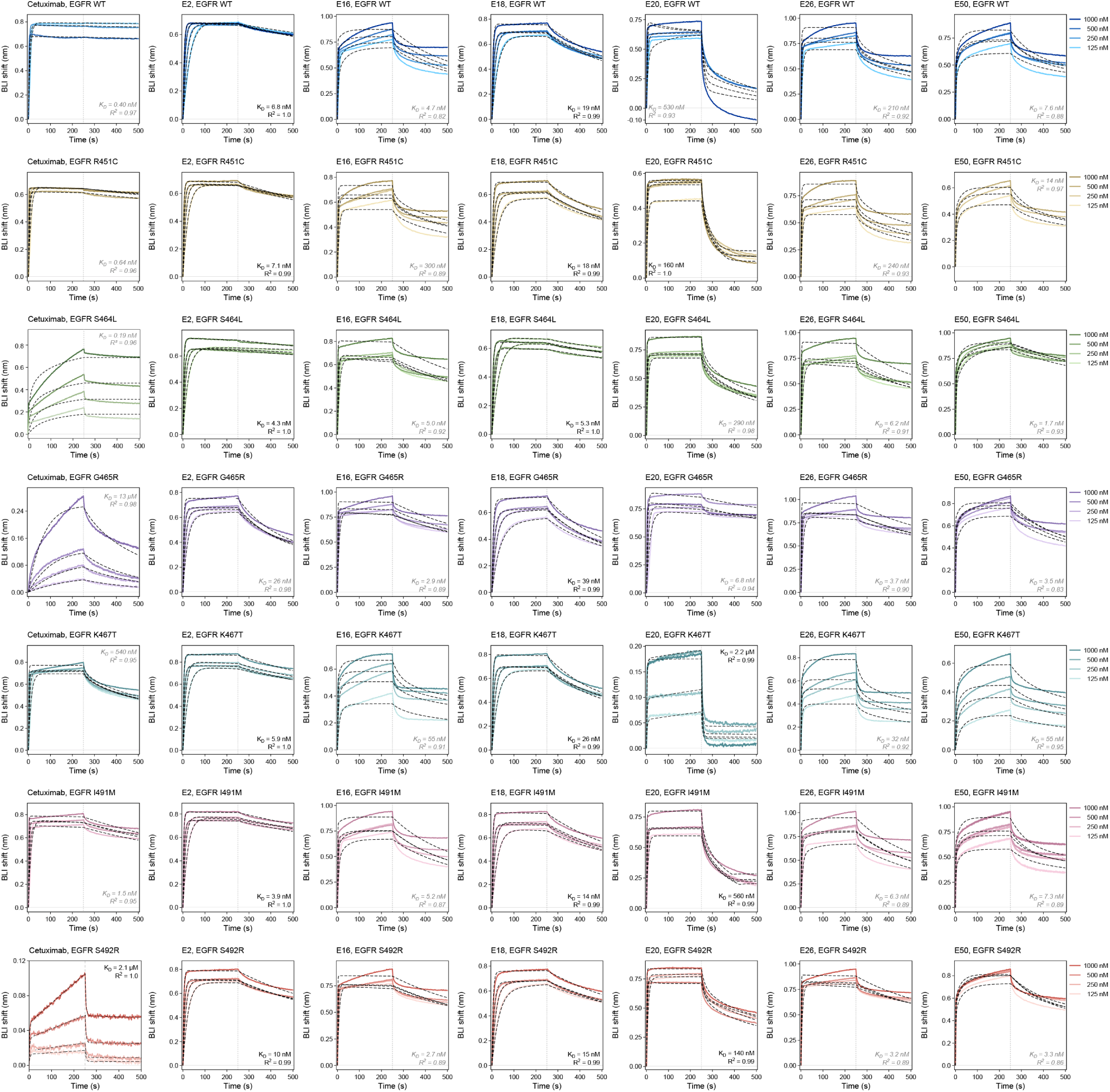
BLI titrations of additional EGFR-binding designs. Concentration-dependent binding of cetuximab, E2, E16, E18, E20, E26, and E50 to wild-type EGFR and the six individual resistance variants. Colored solid traces show reference-subtracted responses at 1,000, 500, 250, and 125 nM antibody; black dashed traces show global fits using the bivalent binding model described in **Methods**. Vertical dotted lines mark the start of dissociation at 250 s. Apparent *K_D_* estimates and corresponding *R*^2^ values are shown where available. Estimates from fits with *R*^2^ < 0.99 are shown in gray italics.

**Figure S5.**
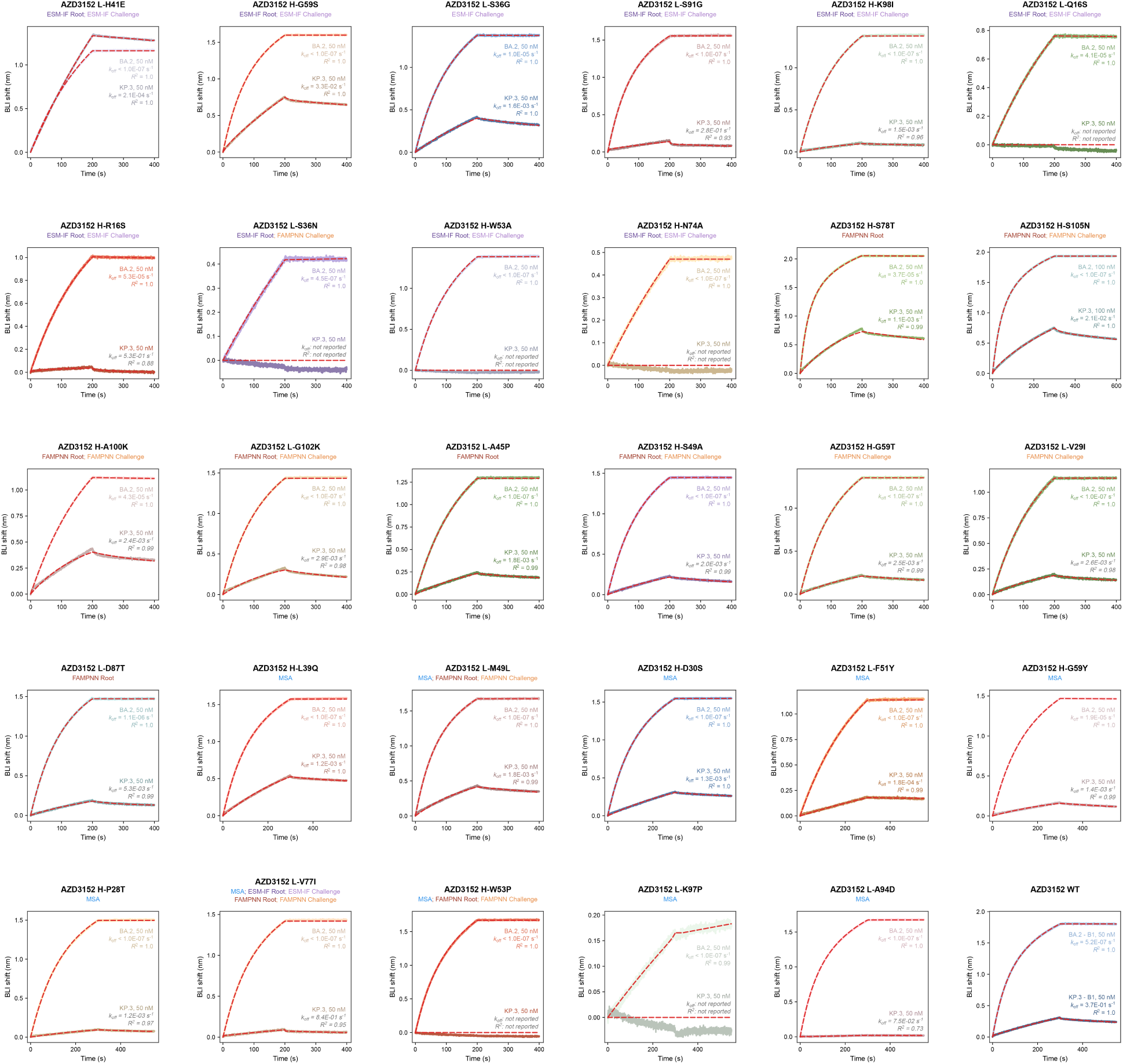
Binding screen of AZD3152 single-substitution variants. BLI sensorgrams for parental AZD3152 and variants carrying individual substitutions proposed using multiple sequence alignments (MSA), ESM-IF or FAMPNN, tested against BA.2 and KP.3 receptor-binding domains (RBDs). Panel titles identify the substitution and proposing method; Root and Challenge denote structural conditioning on BA.2 and KP.3, respectively. H and L denote antibody heavy and light chains. Colored solid traces show reference-subtracted responses, and red dashed traces show kinetic fits where available. Dissociation rate constants (*k_off_*) were estimated from single-concentration screening measurements; corresponding *R*^2^ values are shown. Estimates from fits with *R*^2^<0.99 are shown in gray italics. ‘Not reported’ indicates that the corresponding *k_off_* or *R*^2^ could not be determined.

**Figure S6.**
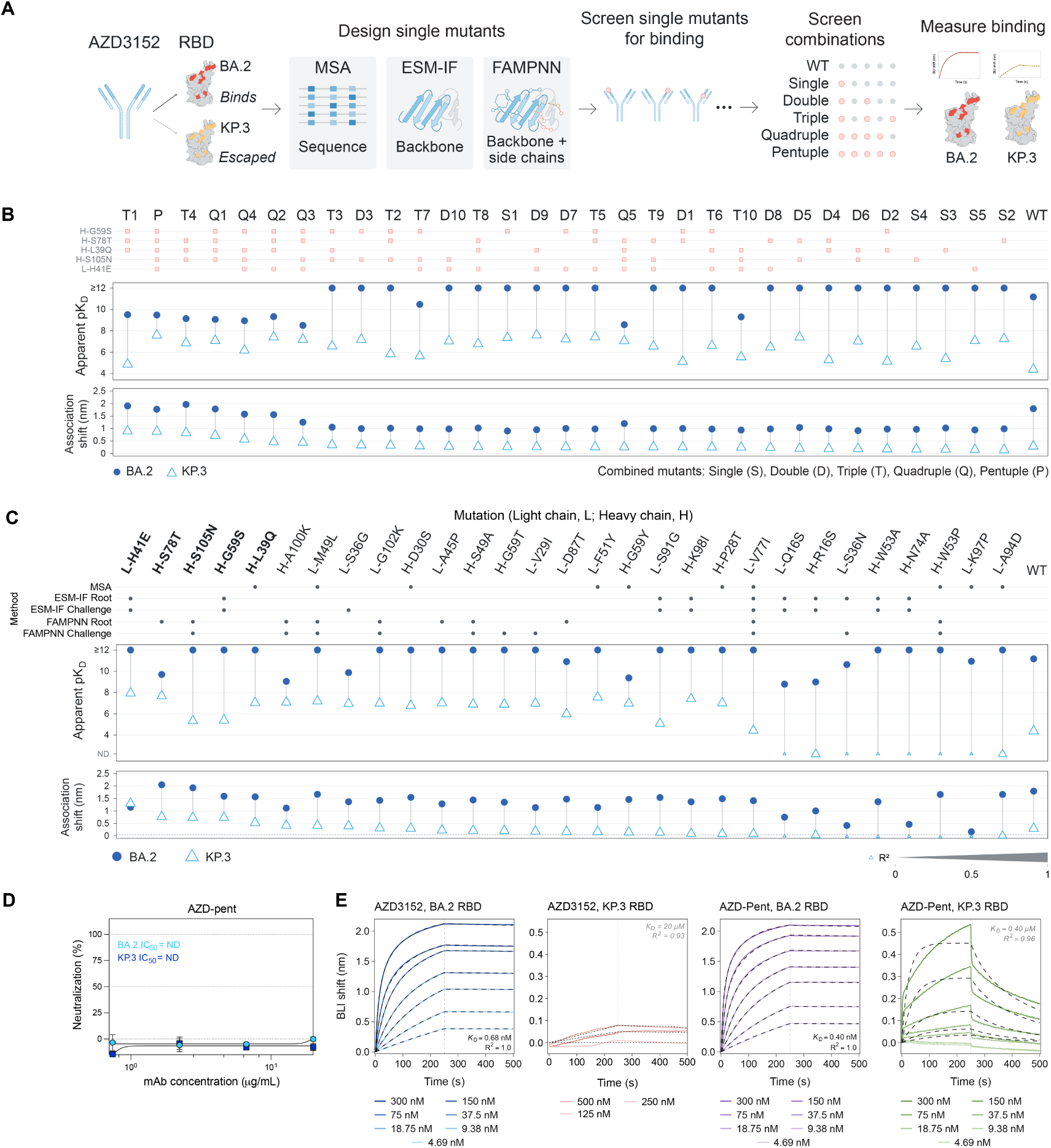
Sequence- and structure-guided benchmarking of AZD3152 rescue. **(A)** Workflow for proposing substitutions using MSA, ESM-IF and FAMPNN and screening individual and combined substitutions against BA.2 and KP.3. **(B)** Apparent *pK_D_* estimates and association responses for selected single-substitution variants and combinations. The matrix shows construct composition; S, D, T, Q and P denote one to five substitutions, respectively, and WT denotes parental AZD3152. **(C)** Initial single-substitution screen, with proposing methods indicated and the five substitutions selected for combination screening in bold. In **(B,C)**, filled circles represent BA.2 and open triangles KP.3; marker size in **(C)** indicates kinetic-fit *R*^2^. Apparent *pK_D_* = − log_10_ [*K_D_*/(M)]. **(D)** Pseudovirus neutralization titrations of the pentuple-substitution variant (AZD-pent) against BA.2 (cyan circles) and KP.3 (blue squares). Despite partial recovery of KP.3 RBD binding, neither variant was detectably neutralized within the tested concentration range. **(E)** BLI titrations of parental AZD3152 and AZD-pent against BA.2 and KP.3 RBDs. Colored solid traces show reference-subtracted responses at the indicated concentrations; black dashed traces show kinetic fits. Vertical dotted lines mark dissociation onset at 250 s. Apparent *K_D_* estimates and *R*^2^ values are annotated; estimates with *R*^2^ < 0.99 appear in gray italics. ND denotes an undetermined estimate. Kinetic fitting is described in **Methods**.

**Figure S7.**
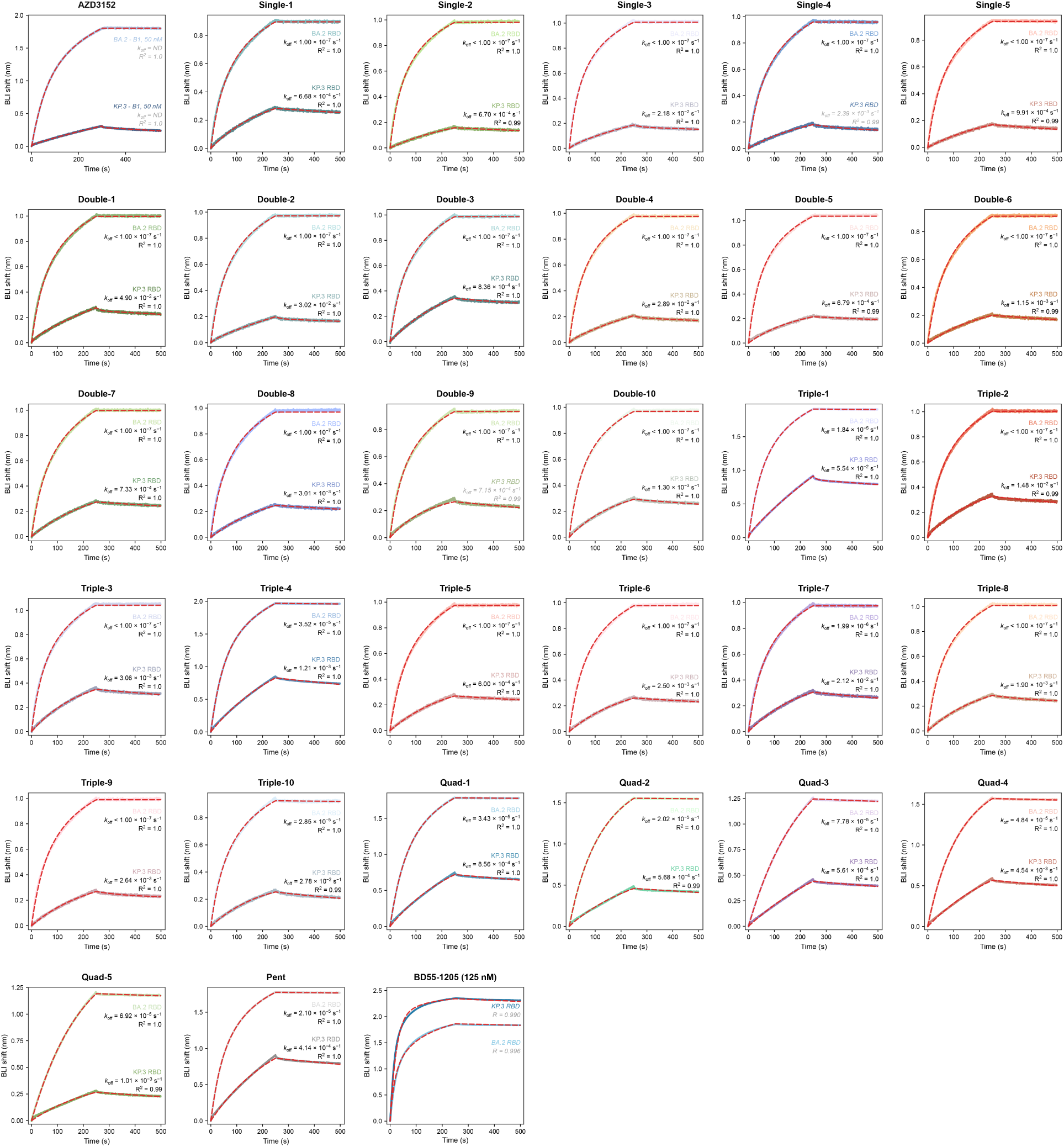
Binding screen of AZD3152 substitution combinations. Single-concentration BLI sensorgrams for the five selected AZD3152 single-substitution variants and their double, triple, quadruple and pentuple combinations against BA.2 and KP.3 RBDs. Construct compositions are shown in **Figure S6B**. Substitution variants and parental AZD3152 were tested at 50 nM antibody; BD55-1205 was included as a positive control at 125 nM. Colored solid traces show reference-subtracted responses, and red dashed traces show kinetic fits where available. Dissociation rate constants (*k_off_*) were estimated from single-concentration screening measurements; corresponding *R*^2^ values are shown. Estimates from fits with *R*^2^<0.99 are shown in gray italics. Blank entries indicate that *k_off_* could not be determined; NR, not reported.

**Figure S8.**
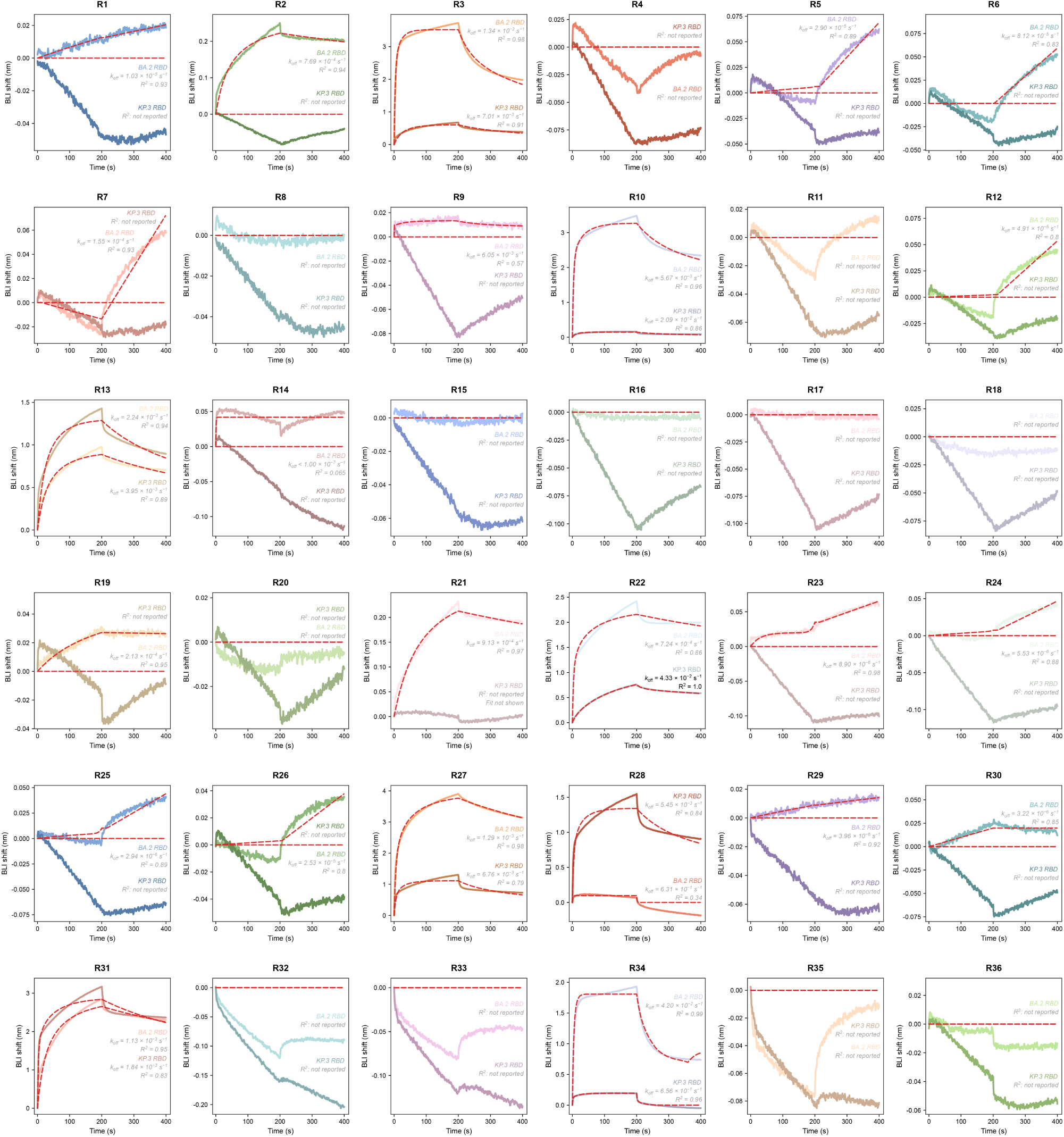
Single-concentration binding screen of BA.2/KP.3 designs R1–R36. BLI sensorgrams for designs R1–R36 tested against BA.2 and KP.3 RBDs at 1,000 nM antibody. Measurements against the two antigens are overlaid and identified by color-matched annotations. Colored solid traces show reference-subtracted responses, and red dashed traces show kinetic fits where available. Dissociation rate constants (*k_off_*) were estimated from single-concentration screening measurements; corresponding *R*^2^ values are shown. Estimates from fits with *R*^2^<0.99 are shown in gray italics. Blank entries indicate that *k_off_* could not be determined; ‘not reported’ indicates that the corresponding *R*^2^ value could not be determined.

**Figure S9.**
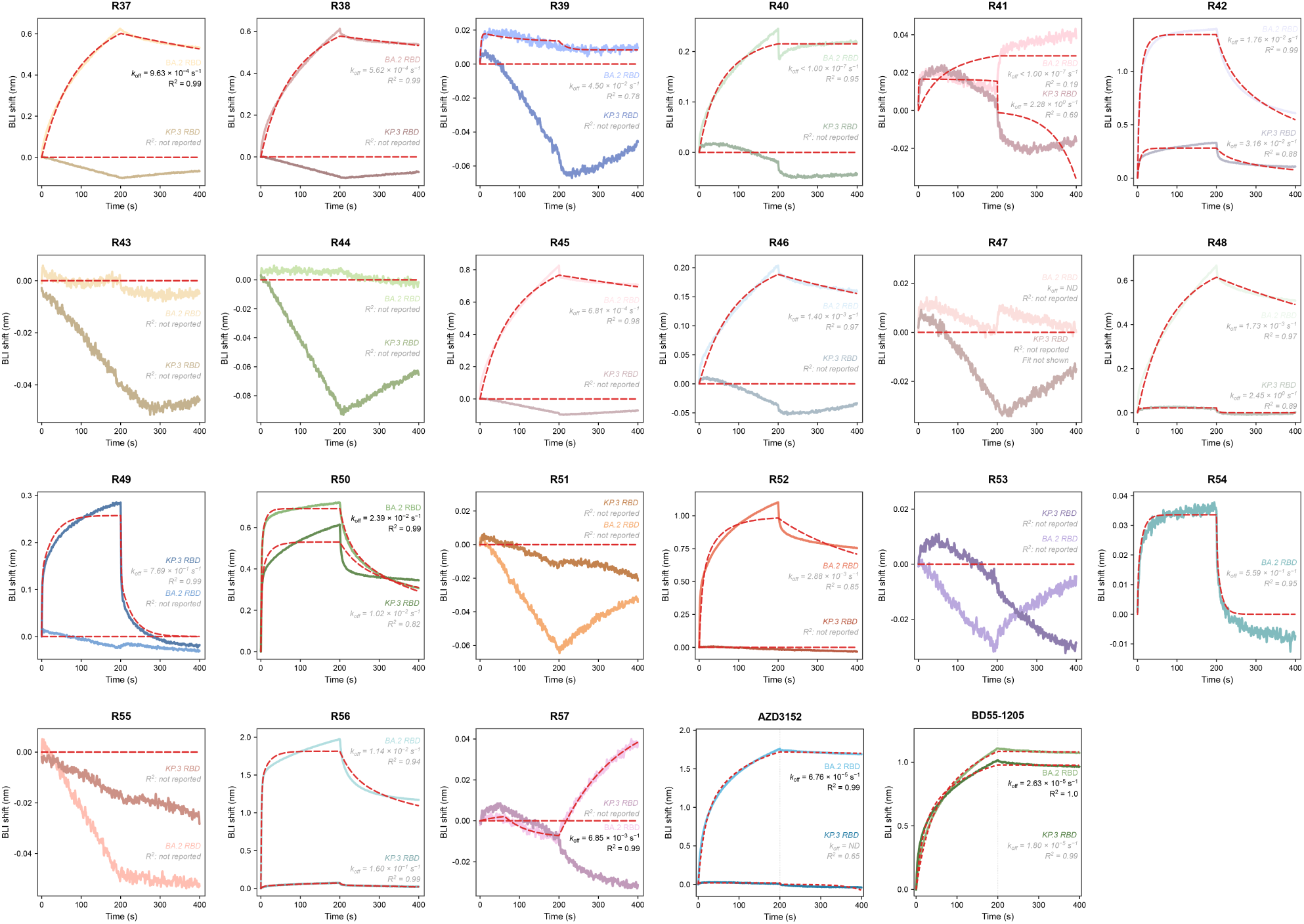
Single-concentration binding screen of BA.2/KP.3 designs R37–R57 and control antibodies. BLI sensorgrams for Germinal Multi designs R37–R57 screened against BA.2 and KP.3 RBDs at 1,000 nM antibody. AZD3152 and BD55-1205 are included as reference controls. Colored solid traces show reference-subtracted responses, and red dashed traces show kinetic fits where available. Dissociation rate constants (*k_off_*) were estimated from single-concentration screening measurements; corresponding *R*^2^ values are shown. Estimates from fits with *R*^2^<0.99 are shown in gray italics. Blank entries indicate that *k_off_* could not be determined; ‘not reported’ indicates that the corresponding *R*^2^ value could not be determined.

**Figure S10.**
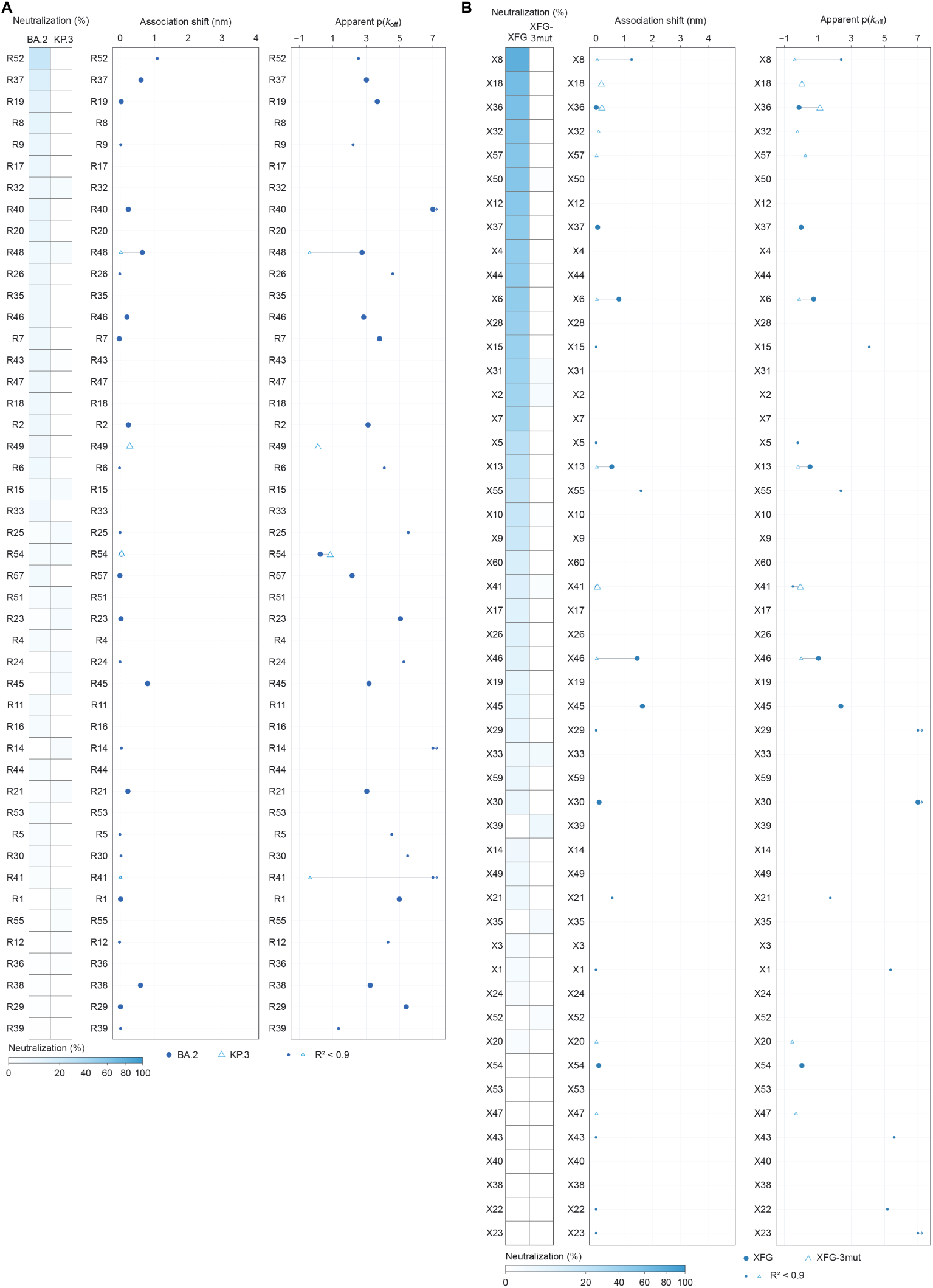
Single-concentration screen of designs not identified as dual-binder leads. Screening results for the remaining designs from the **(A)** BA.2/KP.3 and **(B)** XFG/XFG-Next campaigns, including designs with detectable binding to only one target or no detectable binding to either target under the screening conditions. Each row represents one antibody design. Heatmaps show percent pseudovirus neutralization at an mAb concentration of 10 µg mL^−1^, and adjacent plots show BLI association shifts (nm) and *p*(*k_off_*) values estimated from single-concentration measurements, where *p*(*k*_off)_ = − log_10_ [*k*_off_/(s^−1^)] and *k_off_* is the dissociation rate constant. Filled circles represent the root antigen (BA.2 or XFG), and open triangles represent the challenge antigen (KP.3 or XFG-Next). Lines connect measurements for the same antibody design. Smaller symbols indicate kinetic fits with *R*^2^ < 0.9. ND indicates that a *k*_off_ value could not be determined.

**Figure S11.**
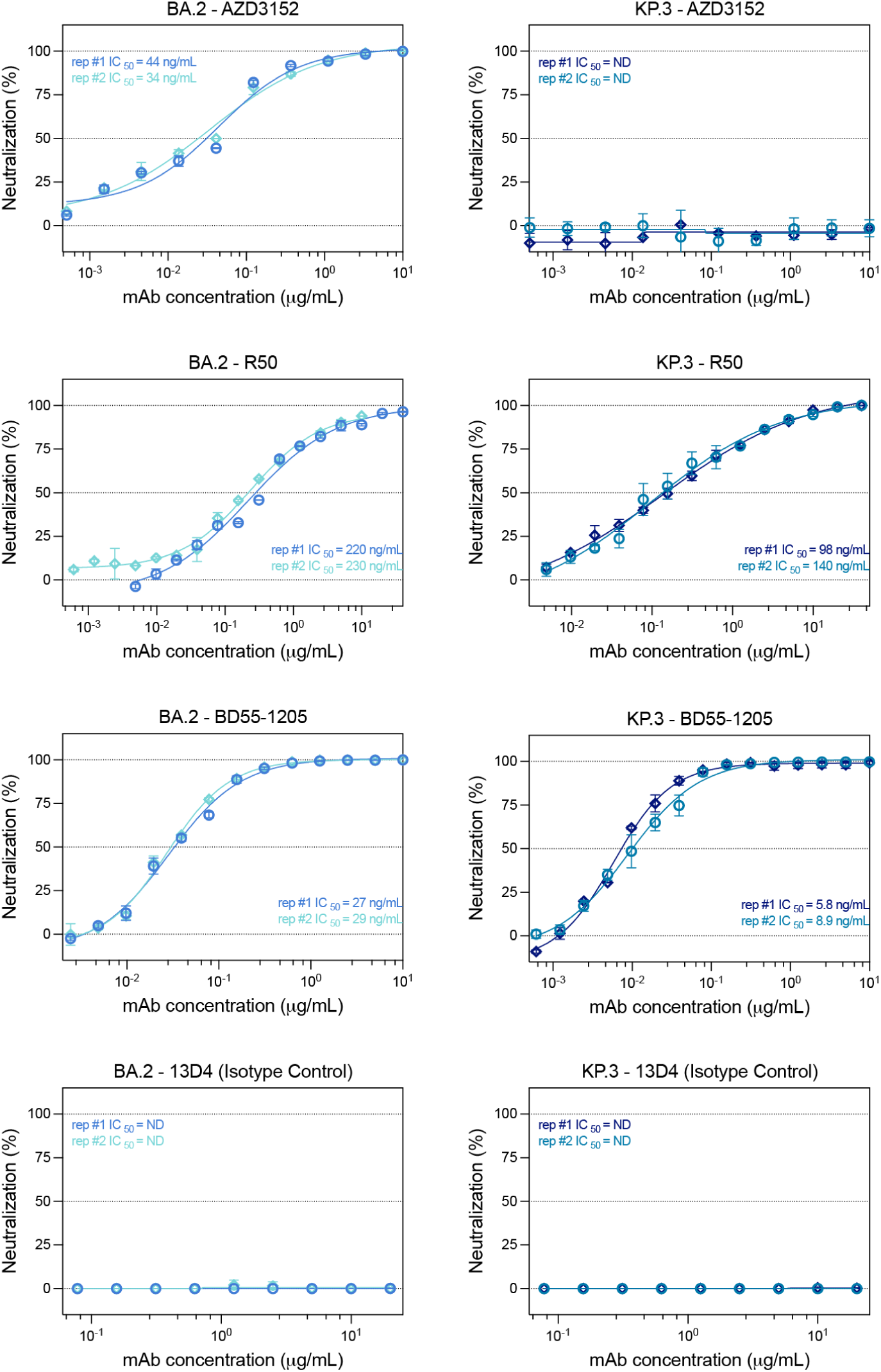
Replicate-resolved neutralization titrations of R50 and reference antibodies against BA.2 and KP.3 pseudoviruses. Neutralization titrations for AZD3152, R50, BD55-1205 and the 13D4 isotype control against BA.2 and KP.3 pseudoviruses. AZD3152 served as a BA.2-active reference antibody, and BD55-1205 served as a positive control for both variants. Two independent experiments, each with two technical replicates per concentration, are shown separately using different colors and symbols. Points show the mean of duplicate measurements within each experiment; error bars indicate SD. Lines show four-parameter logistic fits performed separately for each experiment, with corresponding IC_50_ values annotated in ng/mL. ND indicates that an IC_50_ could not be determined within the tested concentration range.

**Figure S12.**
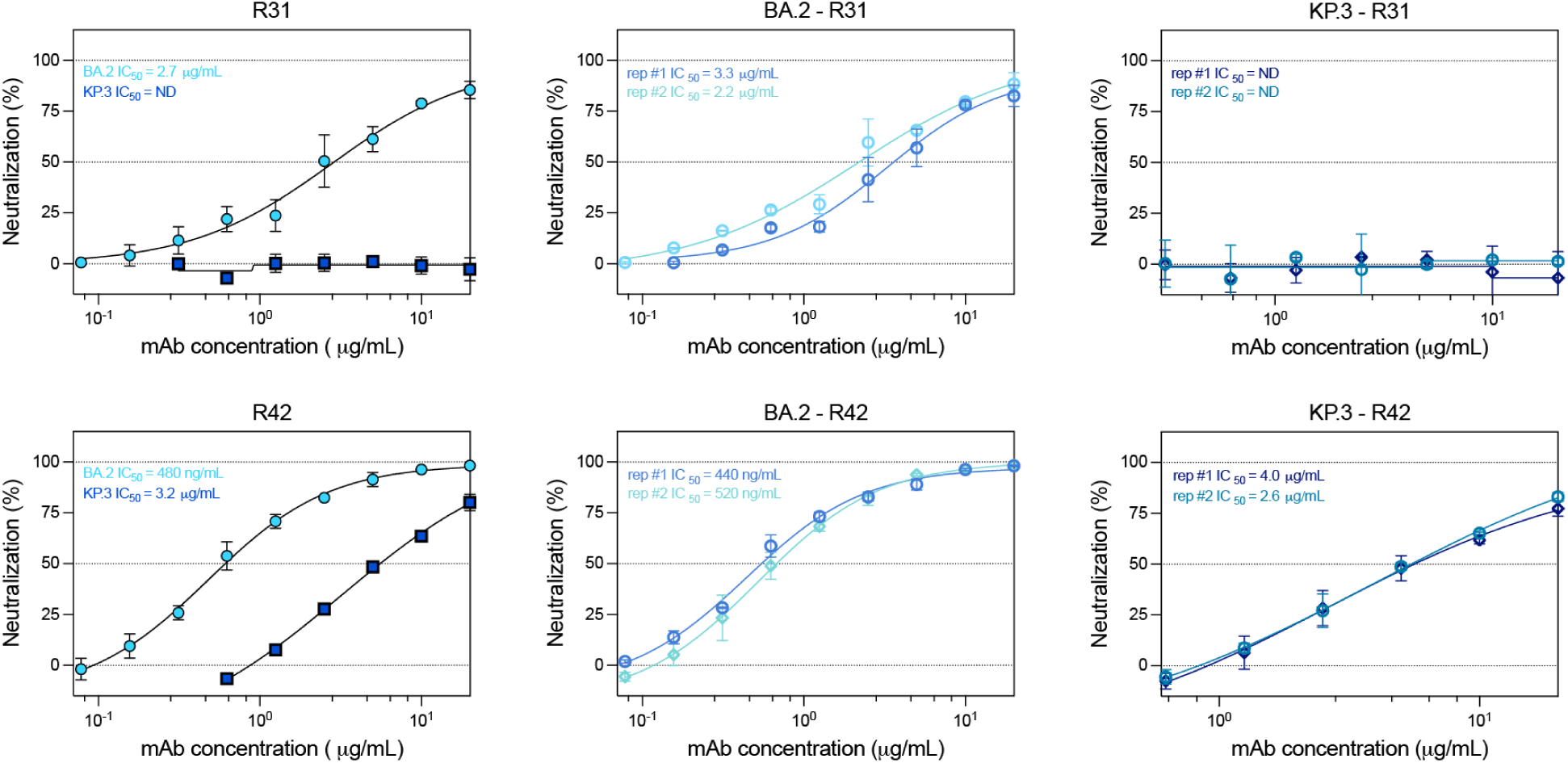
Replicate-resolved neutralization profiles of the dual-binding antibodies R31 and R42 against BA.2 and KP.3. Neutralization titrations of R31 and R42, identified as dual binders by single-point BLI screening. Two independent experiments were performed, each with two technical replicates per concentration. **Left**: comparison of neutralization against BA.2 and KP.3; points show the mean of the two experiment-level means, and error bars indicate SD. Annotated IC_50_ values are the geometric means of the unrounded fitted estimates from two independent experiments, expressed in ng/mL and reported to two significant figures. **Middle** and **right**: the experiments are shown separately for BA.2 and KP.3, respectively. Points show the mean of duplicate measurements within each experiment; error bars indicate SD. Lines show four-parameter logistic fits performed separately for each experiment, with corresponding IC_50_ values expressed in *μ*g/mL or ng/mL, as indicated. ND indicates that an IC_50_ could not be determined within the tested concentration range.

**Figure S13.**
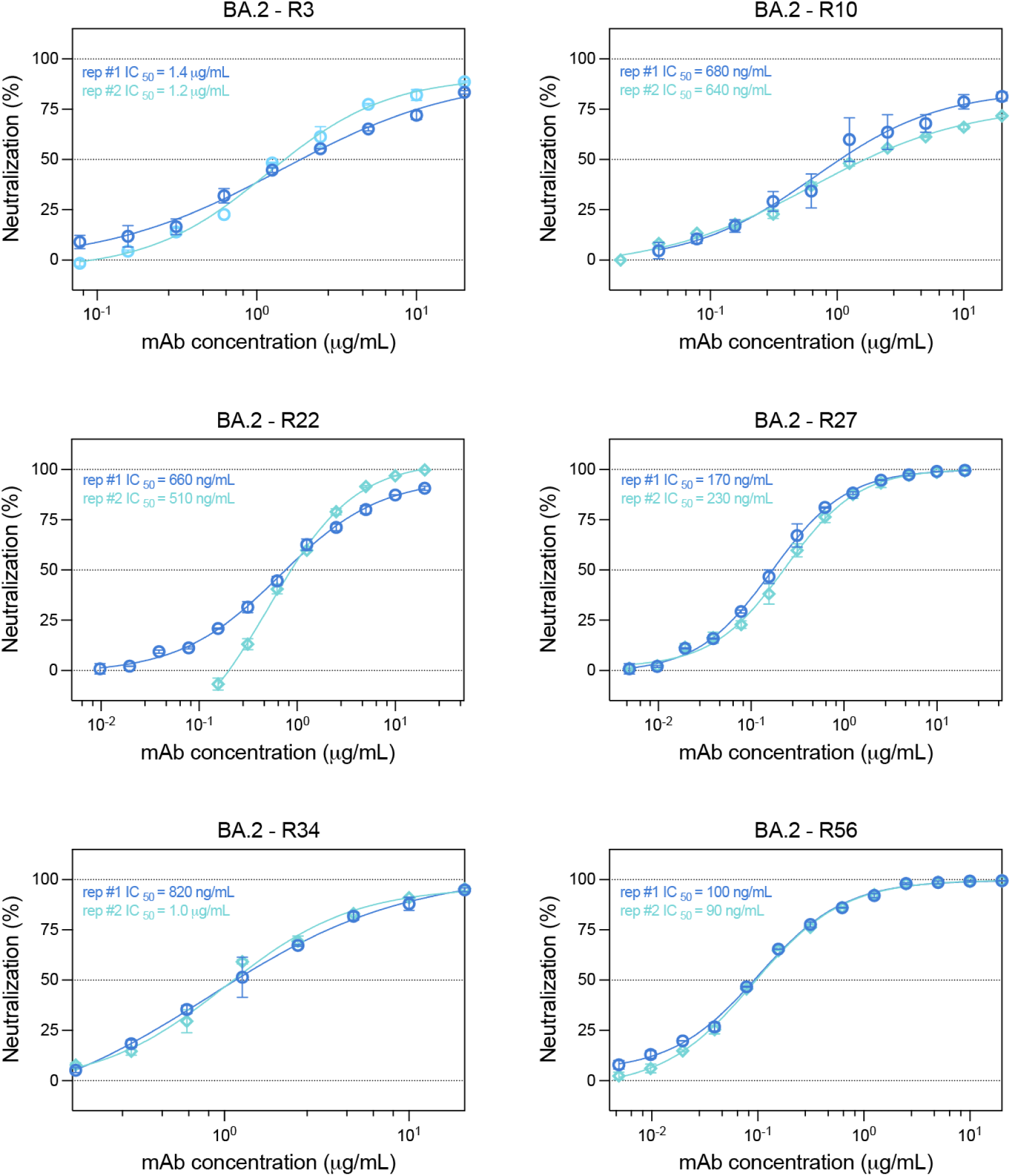
Replicate-resolved neutralization titrations of designs classified as BA.2-only neutralizers. BA.2 neutralization curves for R3, R10, R22, R27, R34 and R56, identified through single-point neutralization and BLI binding screening; dual neutralizers are excluded. Each curve represents one of two independent experiments. Points show the mean of two technical replicates within each experiment; error bars indicate SD. Lines show four-parameter logistic fits performed separately for each experiment, with corresponding IC_50_ values expressed in *μ*g/mL or ng/mL, as indicated.

**Figure S14.**
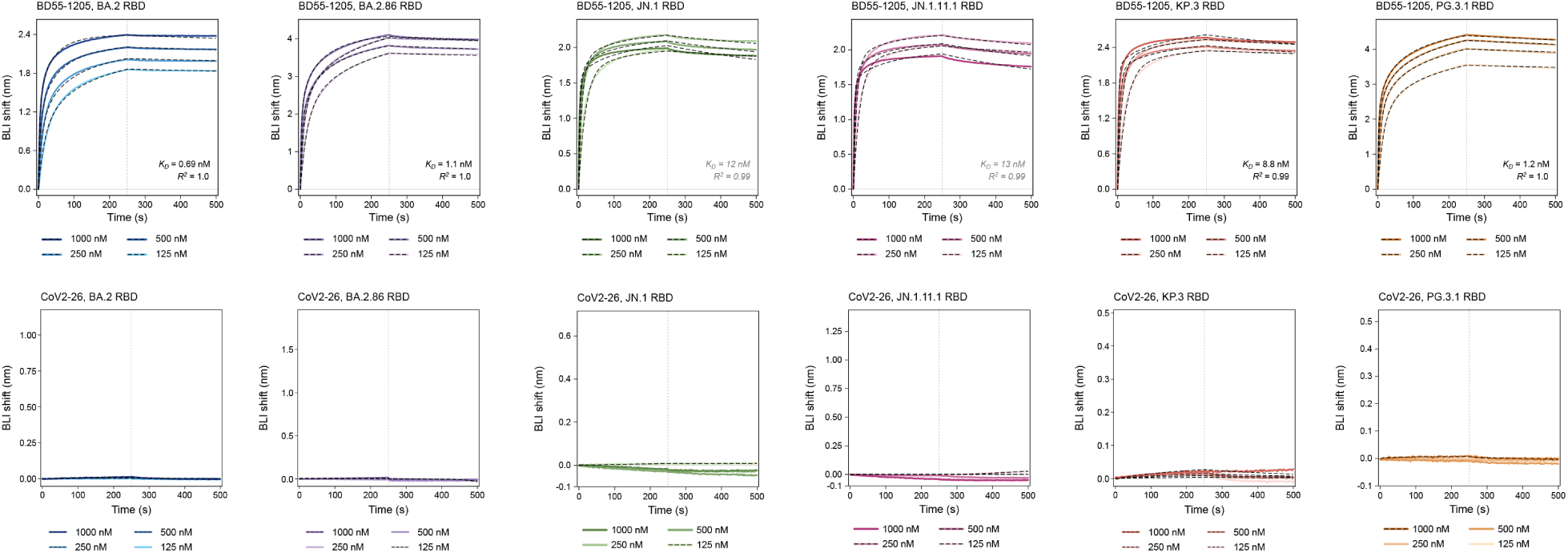
Reference-antibody binding across the SARS-CoV-2 RBD variant panel. BLI titrations of BD55-1205 (upper row) and CoV2-26 (lower row), included as positive and negative binding controls, respectively, against BA.2, BA.2.86, JN.1, JN.1.11.1, KP.3 and PG.3.1 RBDs. Colored solid traces show reference-subtracted responses at the indicated antibody concentrations; black dashed traces show kinetic fits where available. Vertical dotted lines mark the start of dissociation at 250 s. Apparent *K_D_* estimates and corresponding *R*^2^ values are shown where available. Estimates from fits with *R*^2^ < 0.99 are shown in gray italics. Kinetic fitting is described in **Methods**.

**Figure S15.**
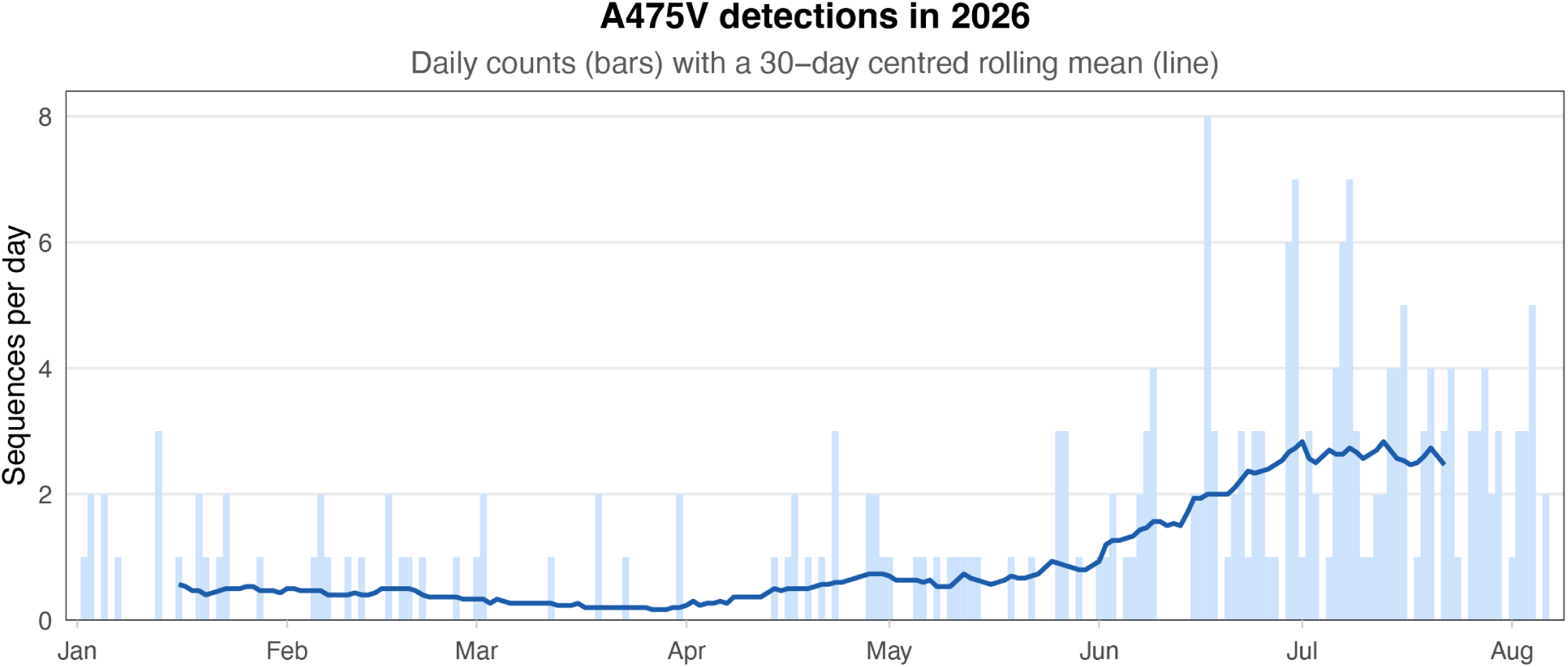
Detection counts of the spike A475V substitution in SARS-CoV-2 sequence data. Daily counts of sequences carrying A475V in the analyzed dataset from January through August 2026. Pale blue bars show daily sequence counts; the dark blue line shows the 30-day centered rolling mean. Counts represent absolute detections rather than substitution frequencies.

**Figure S16.**
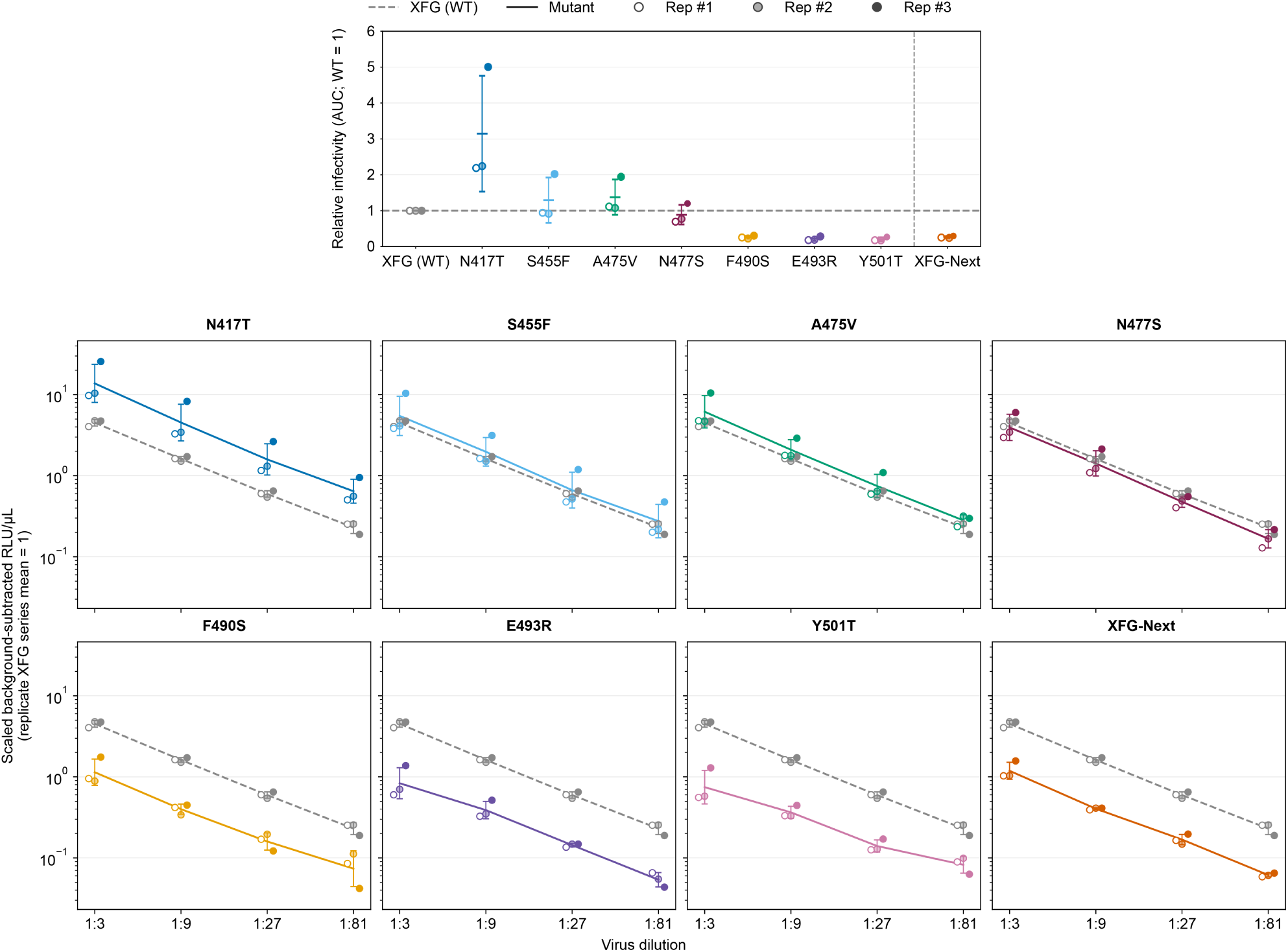
Pseudovirus entry measurements for XFG spike variants. **Upper panel**: area under the dilution-response curve (AUC) for each indicated XFG variant, normalized to the matched parental XFG value. **Lower panels**: background-subtracted luminescence per unit volume (RLU/µL) across the indicated pseudovirus dilution series. Within each replicate, responses are scaled so that the mean of the parental XFG dilution series equals 1. Colored solid lines show the indicated variants; gray dashed lines show parental XFG. Symbol fill distinguishes the three plotted replicate series. In the upper panel, summary points show the mean of the three independent measurements; error bars indicate SD (*n* = 3). Preparations were not particle-normalized; these measurements therefore demonstrate detectable spike-mediated entry rather than differences in intrinsic infectivity.

**Figure S17.**
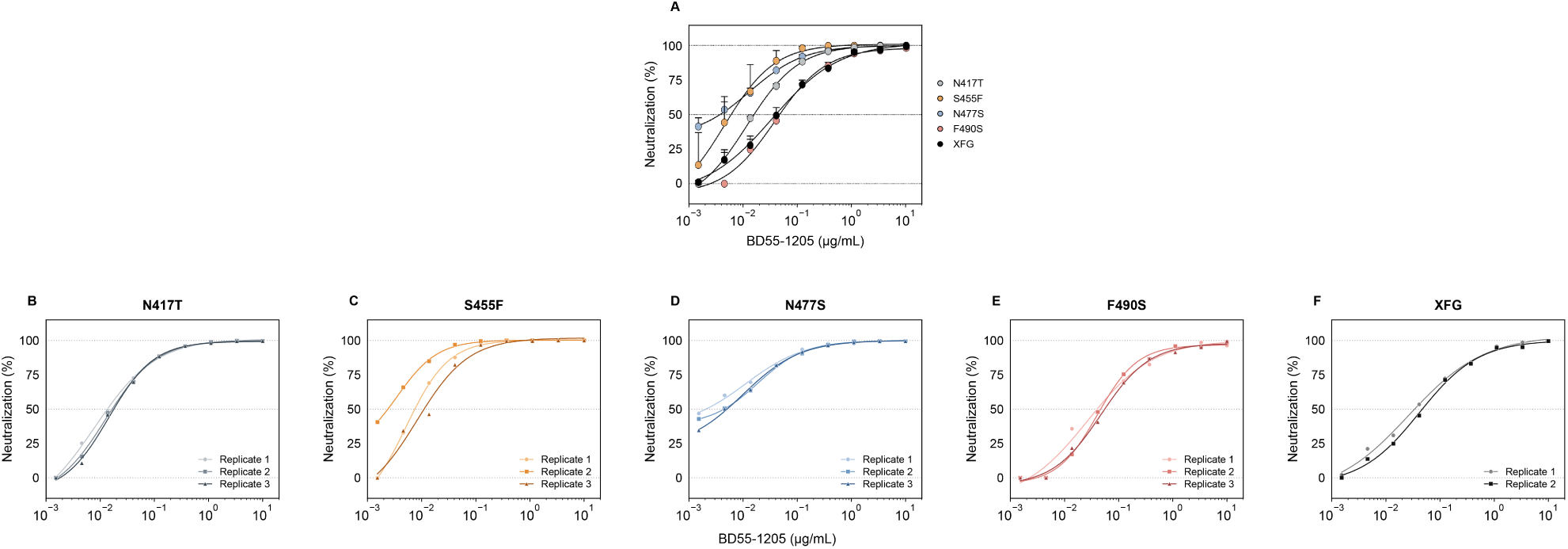
BD55-1205 neutralization of XFG variants carrying antiGen-predicted substitutions outside the six escape-associated sites. **(A)** Concentration-dependent neutralization of parental XFG and the indicated single-substitution variants. Points show the mean across replicates; error bars indicate SD, and solid lines show fitted concentration-response curves. **(B–F)** Individual replicate measurements for N417T, S455F, N477S, F490S and parental XFG, respectively. Colors and marker shapes distinguish replicates. Solid lines represent four-parameter logistic fits. Measurements comprise three replicates per mutant and two for parental XFG.

**Figure S18.**
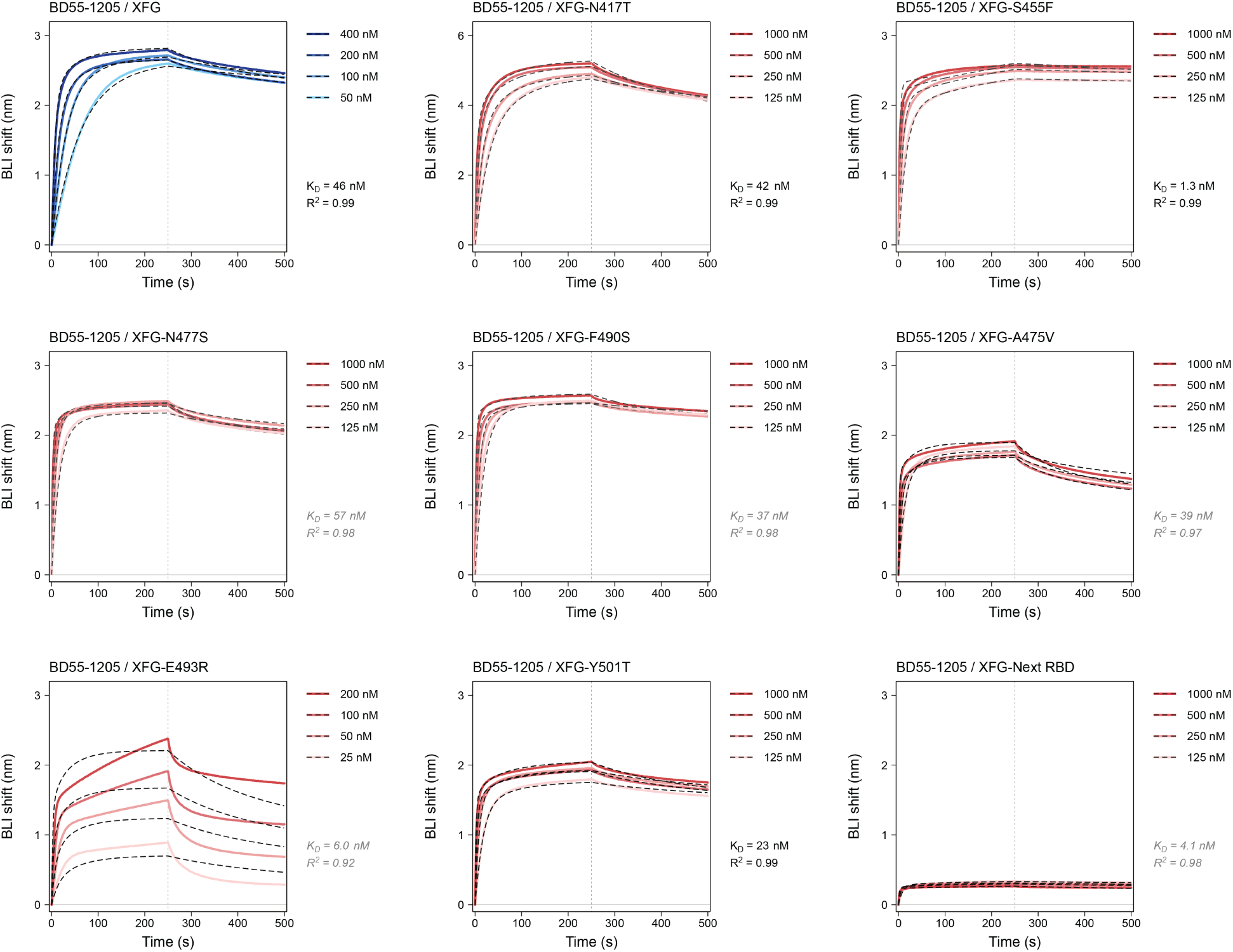
BD55-1205 binding to antiGen-predicted XFG RBD variants. BLI titrations of BD55-1205 against parental XFG, the indicated single-substitution RBD variants (N417T, S455F, A475V, N477S, F490S, E493R and Y501T), and the combined XFG-Next challenge antigen. Blue traces show binding to parental XFG; coral traces show binding to the variant RBDs at the antibody concentrations specified in each panel. Solid traces show reference-subtracted responses; black dashed traces show kinetic fits. Vertical dotted lines mark the start of dissociation at 250 s. Apparent *K_D_* estimates and corresponding *R*^2^ values are annotated. Estimates from fits with *R*^2^<0.99 are shown in gray italics.Kinetic fitting is described in **Methods**.

**Figure S19.**
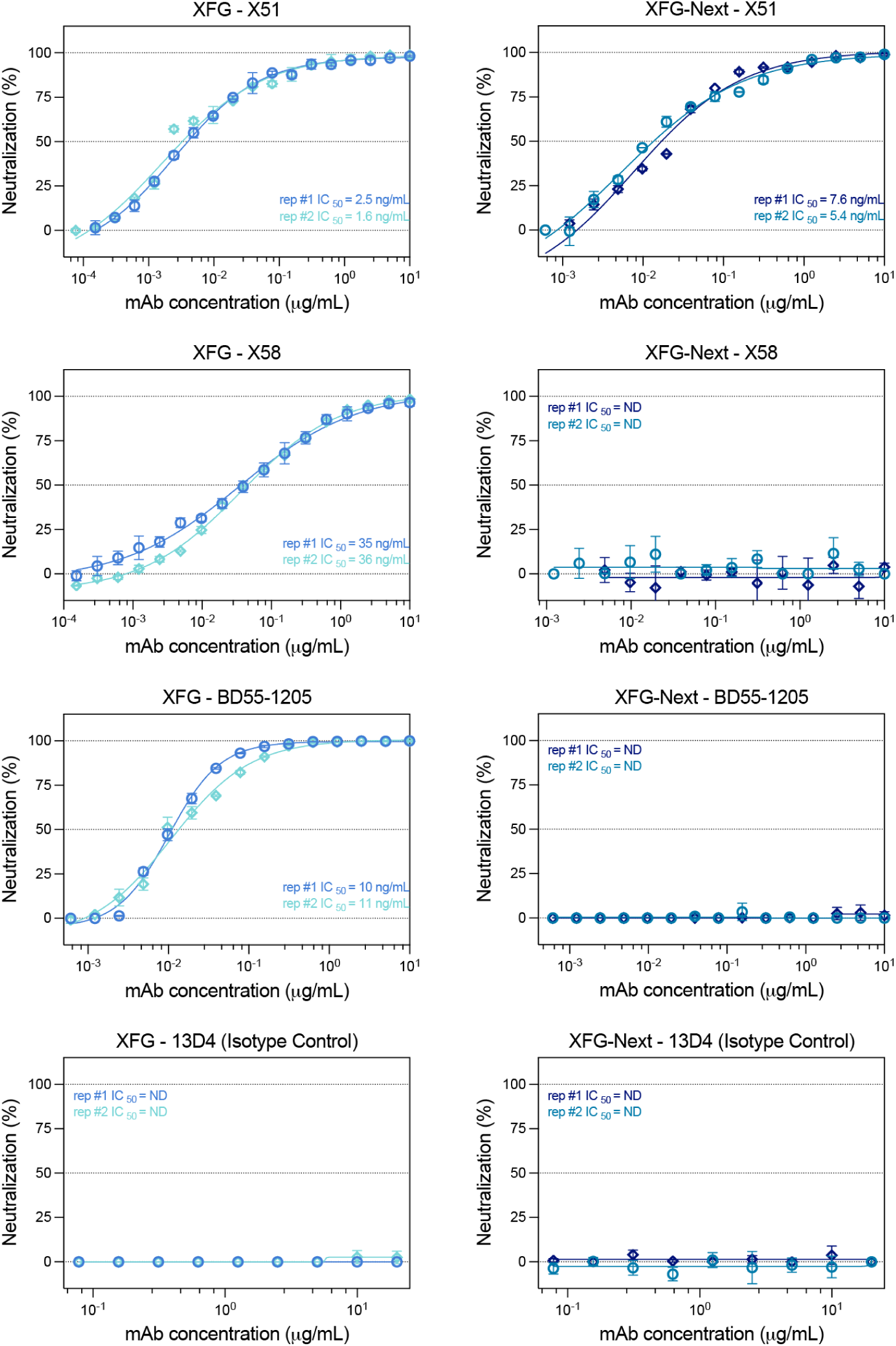
Replicate-resolved neutralization titrations of X51, X58 and reference antibodies against XFG and XFG-Next pseudoviruses. Neutralization titrations for X51, X58, parental BD55-1205 and the 13D4 isotype control against XFG and XFG-Next (A475V/E493R/Y501T) pseudoviruses. Two independent experiments, each with two technical replicates per concentration, are shown separately using different colors and symbols. Points show the mean of duplicate measurements within each experiment; error bars indicate SD. Lines show four-parameter logistic fits performed separately for each experiment, with corresponding IC_50_ values annotated in ng/mL. ND indicates that an IC_50_ could not be determined within the tested concentration range.

**Figure S20.**
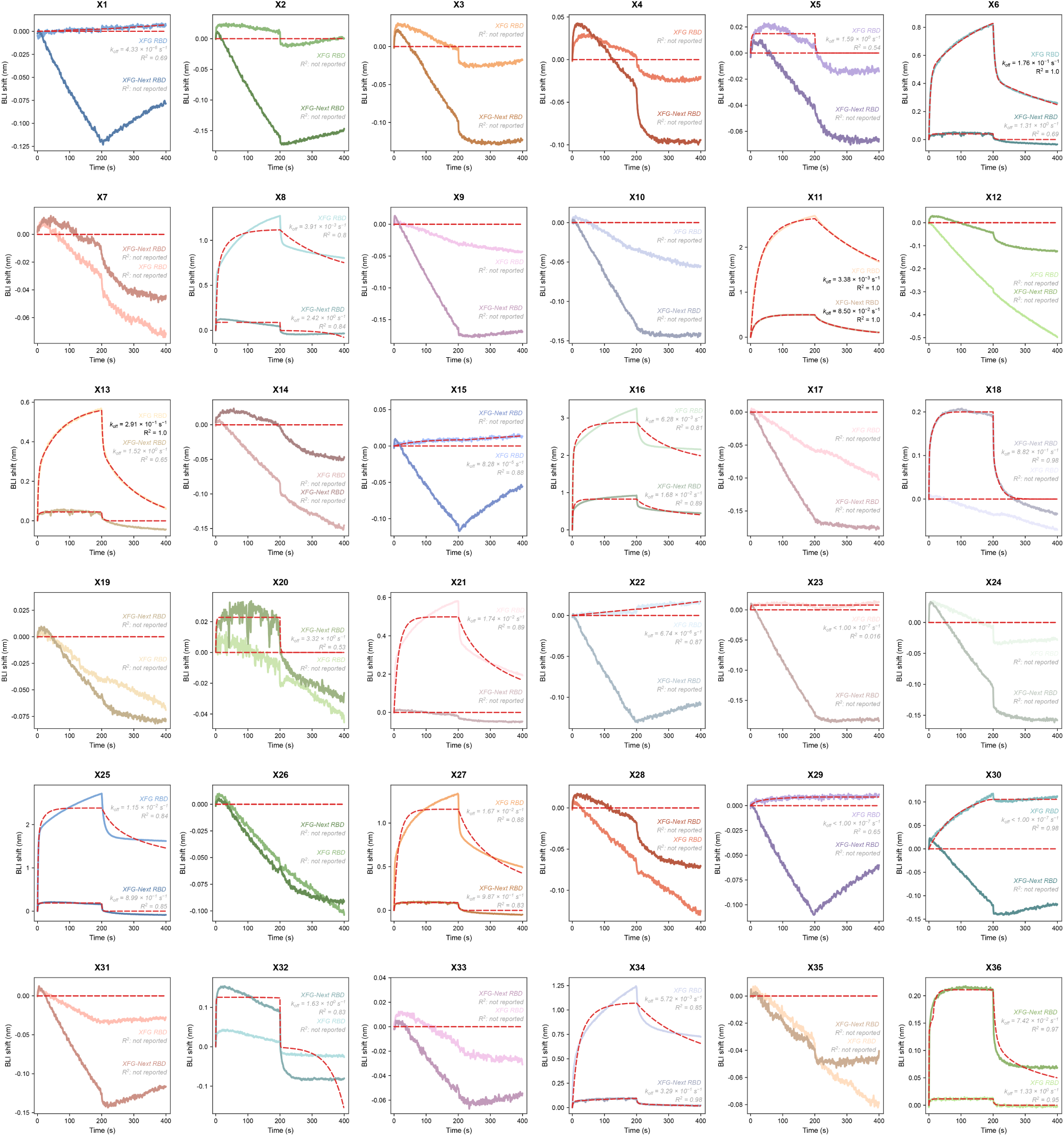
Single-concentration binding screen of XFG/XFG-Next designs X1–X36. BLI sensorgrams for BD55-1205-derived designs X1–X36 tested against parental XFG and XFG-Next RBDs at 1,000 nM antibody. Measurements against the two antigens are overlaid and identified by color-matched annotations. Colored solid traces show reference-subtracted responses, and red dashed traces show kinetic fits where available. Dissociation rate constants (*k_off_*) were estimated from single-concentration screening measurements; corresponding *R*^2^ values are shown. Estimates from fits with *R*^2^ < 0.99 are shown in gray italics. Blank entries indicate that *k_off_* could not be determined; ‘not reported’ indicates that the corresponding *R*^2^ value could not be determined.

**Figure S21.**
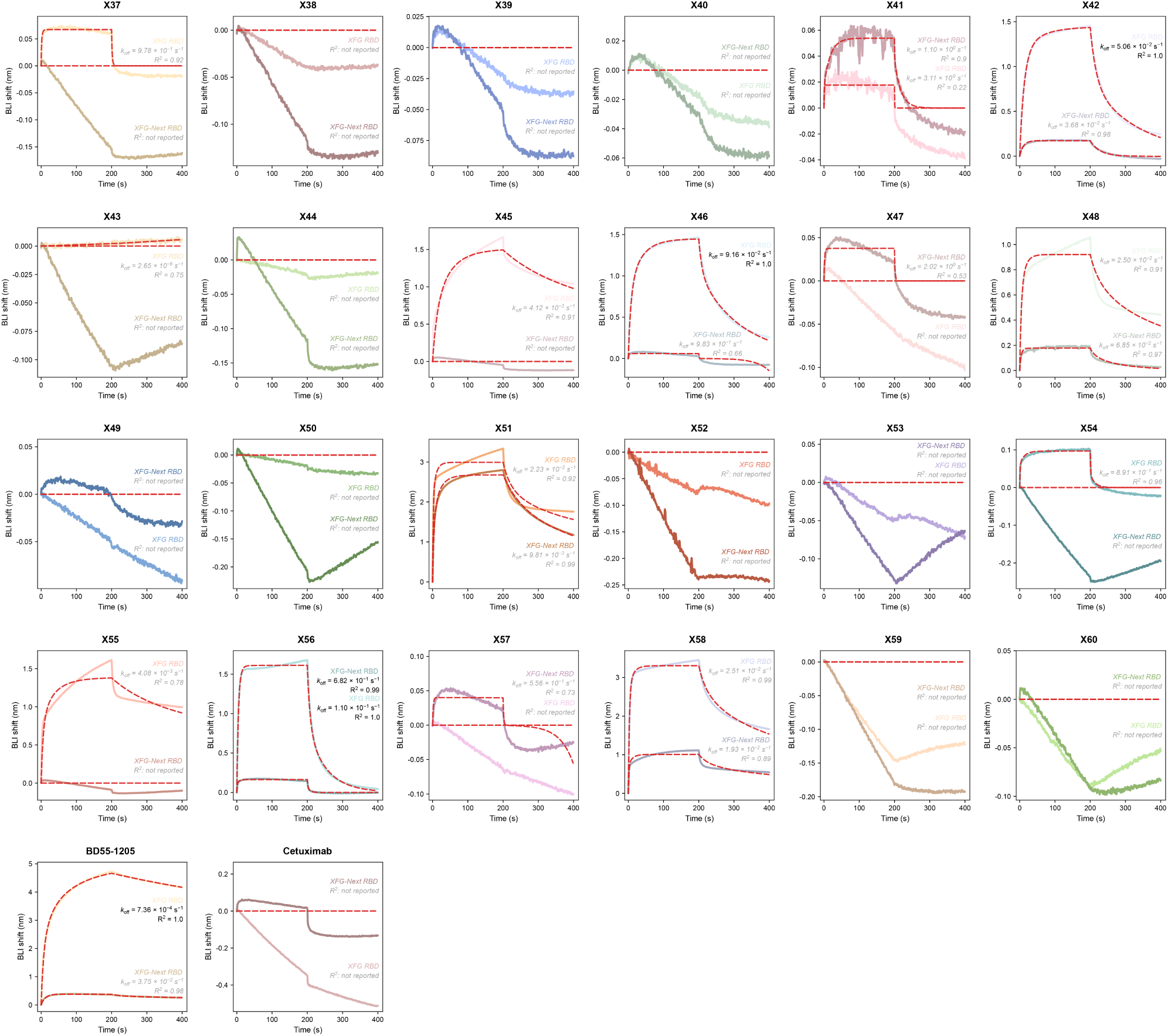
Single-concentration binding screen of XFG/XFG-Next designs X37–X60 and controls. BLI sensorgrams for BD55-1205-derived designs X37–X60 screened against XFG and XFG-Next RBDs at 1,000 nM antibody. Parental BD55-1205 and the unrelated antibody cetuximab are included as controls. Measurements against the two antigens are identified by color-matched annotations. Colored solid traces show reference-subtracted responses, and red dashed traces show kinetic fits where available. Dissociation rate constants (*k_off_*) were estimated from single-concentration screening measurements; corresponding *R*^2^ values are shown. Estimates from fits with *R*^2^ < 0.99 are shown in gray italics. Blank entries indicate that *k_off_* could not be determined; ‘not reported’ indicates that the corresponding *R*^2^ value could not be determined.

**Figure S22.**
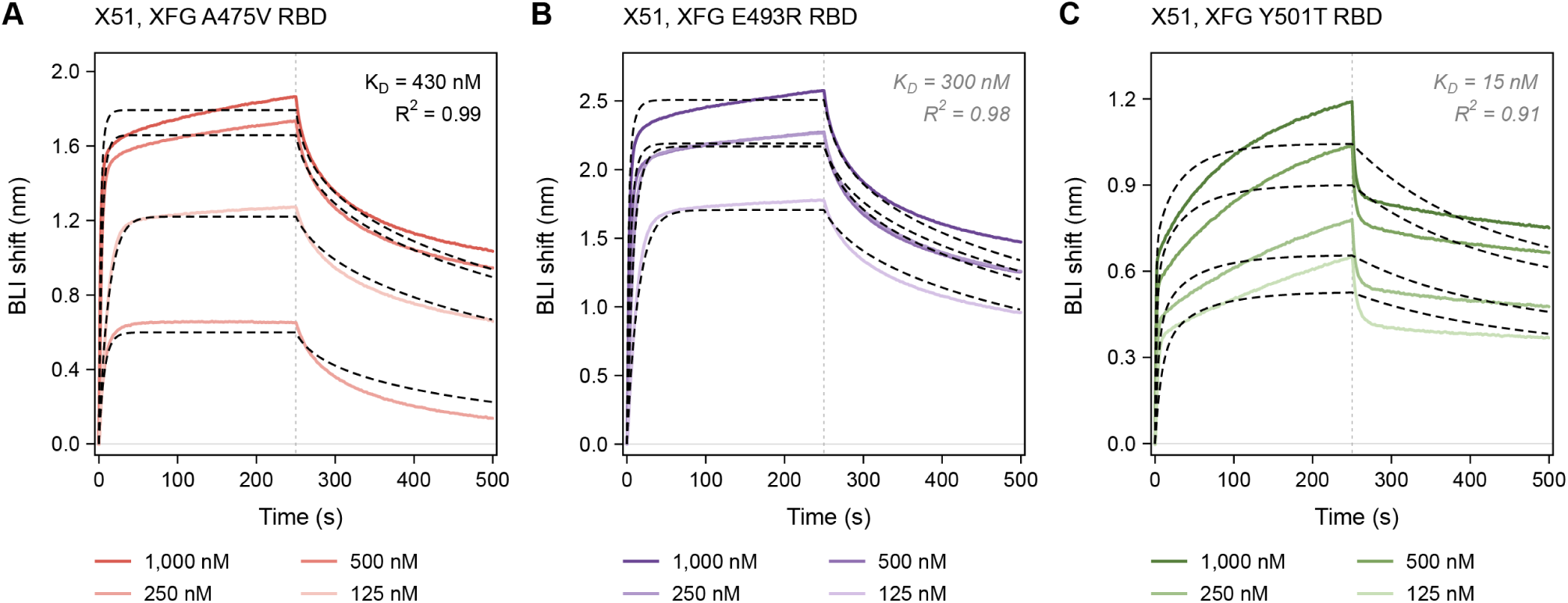
X51 binding to individual XFG challenge variants. BLI titrations of X51 against the individual XFG RBD variants labelled **(A)** A475V, **(B)** E493R and **(C)** Y501T. Colored solid traces show reference-subtracted responses at the indicated antibody concentrations; black dashed traces show global fits using the bivalent binding model described in **Methods**. Vertical dotted lines mark the start of dissociation at 250 s. Apparent *K_D_* and corresponding *R*^2^ values are annotated. Gray italic annotations denote displayed *R*^2^ < 0.99.

**Figure S23.**
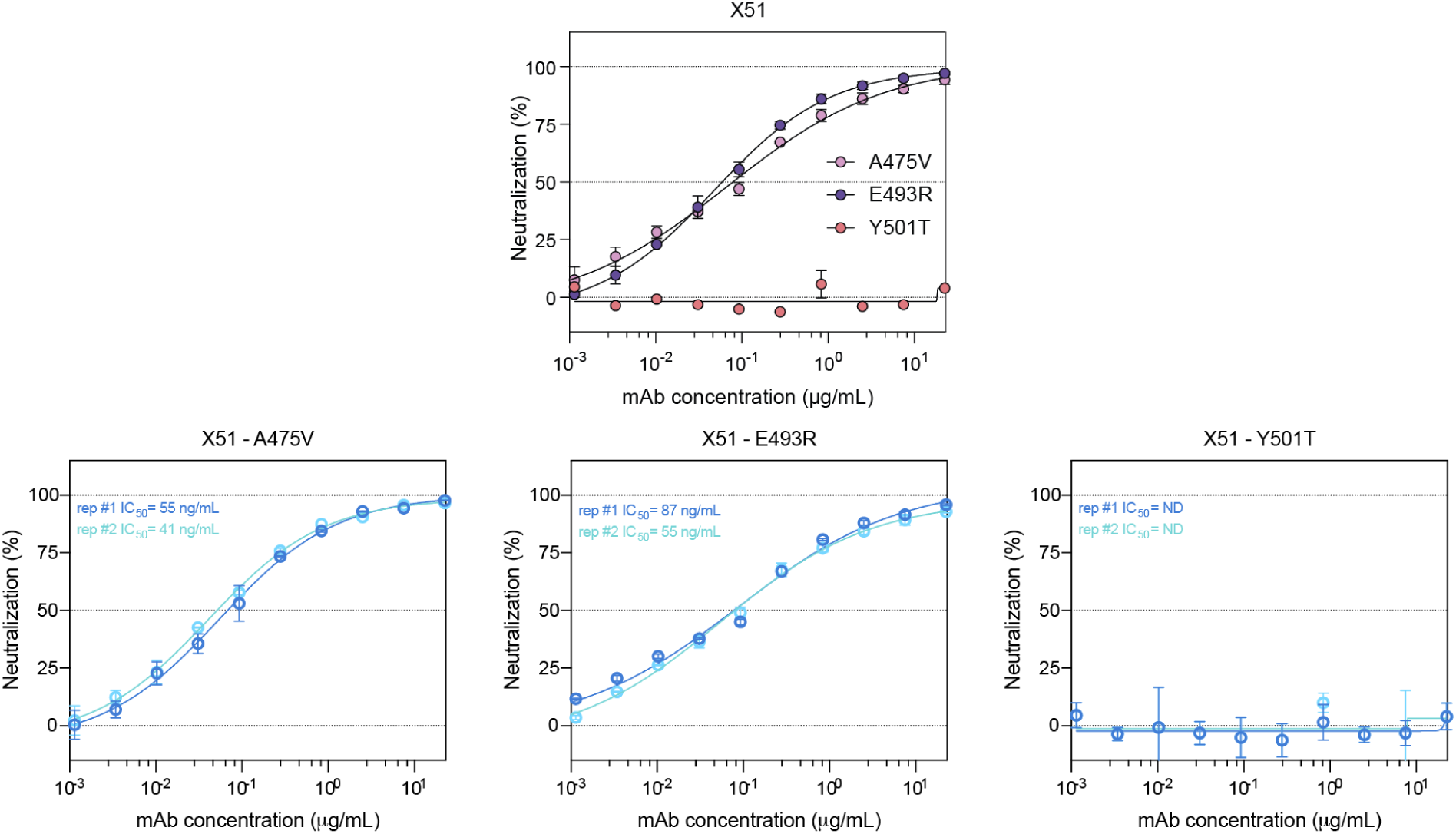
X51 neutralization of XFG variants carrying the individual substitutions comprising XFG-Next. Neutralization titrations of X51 against XFG pseudoviruses carrying the individual A475V, E493R or Y501T substitution. Two independent experiments were performed, each with two technical replicates per concentration. **Top**: comparison of the three variants; points show the mean of the two experiment-level means, and error bars indicate SD. **Bottom**: the experiments are shown separately for each variant using two tones of the corresponding color. Points show the mean of duplicate measurements within each experiment; error bars indicate SD. Lines show four-parameter logistic fits performed separately for each experiment, with corresponding IC_50_ values annotated in ng/mL. ND indicates that an IC_50_ could not be determined within the tested concentration range.

**Figure S24.**
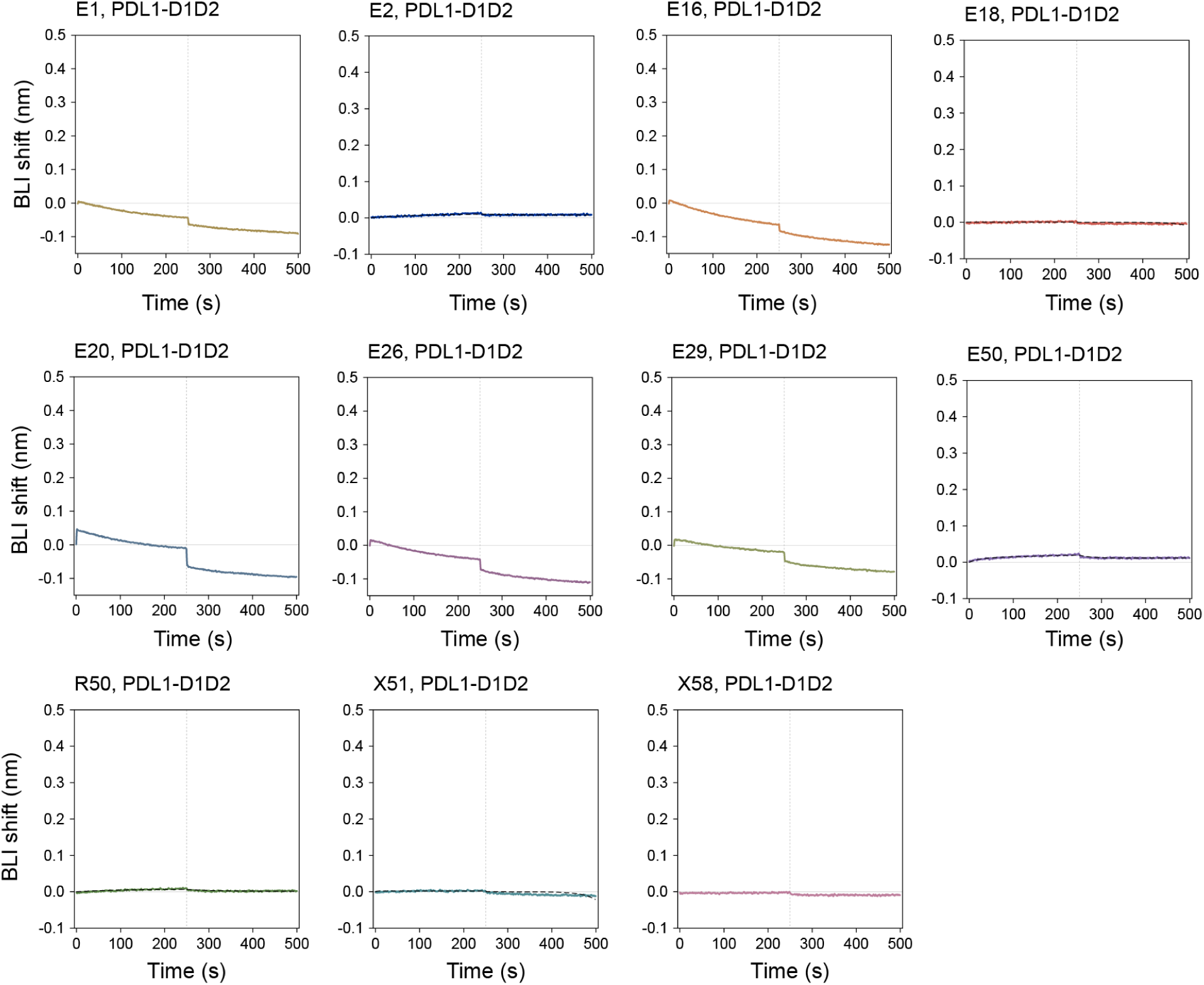
Off-target BLI screening of selected designs against the unrelated antigen PD-L1. Single-concentration BLI sensorgrams for E1, E2, E16, E18, E20, E26, E29, E50, R50, X51 and X58 tested against the PD-L1 D1–D2 extracellular construct at 1,000 nM antibody. Colored traces show reference-subtracted responses; vertical dotted lines mark the start of dissociation at 250 s. No appreciable positive association response is evident under the conditions tested.

**Figure S25.**
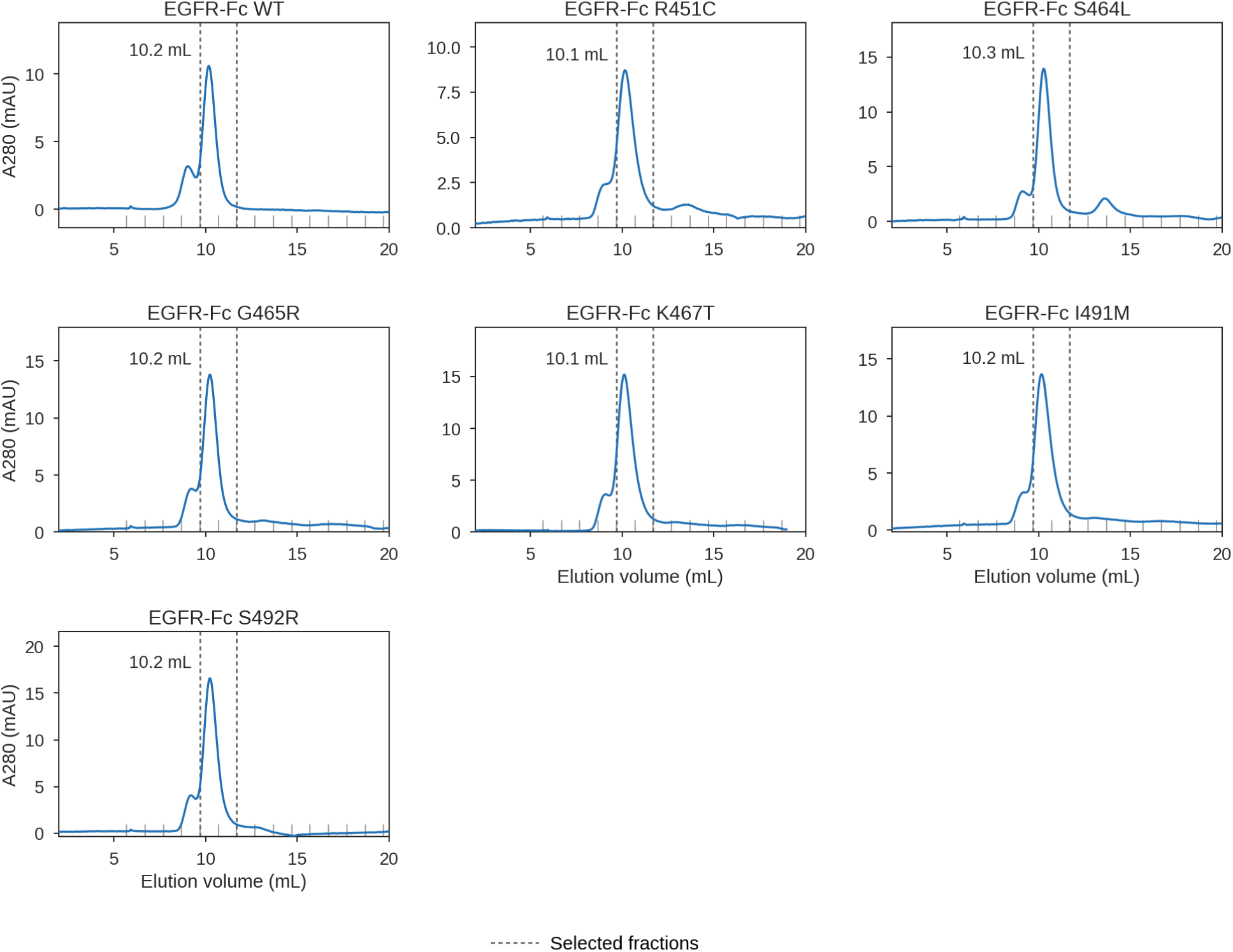
Size-exclusion chromatography of the EGFR-Fc antigen panel. Size-exclusion chromatograms for wild-type EGFR-Fc and the individual R451C, S464L, G465R, K467T, I491M and S492R variants. Traces show absorbance at 280 nm as a function of elution volume. Gray baseline ticks indicate collected fraction boundaries at 1 mL intervals; dashed vertical lines delimit the fractions pooled for downstream experiments (fractions 6–7, 9.7–11.7 mL). Annotated values indicate the elution volume of the principal peak within the selected 8–12 mL product region.

**Figure S26.**
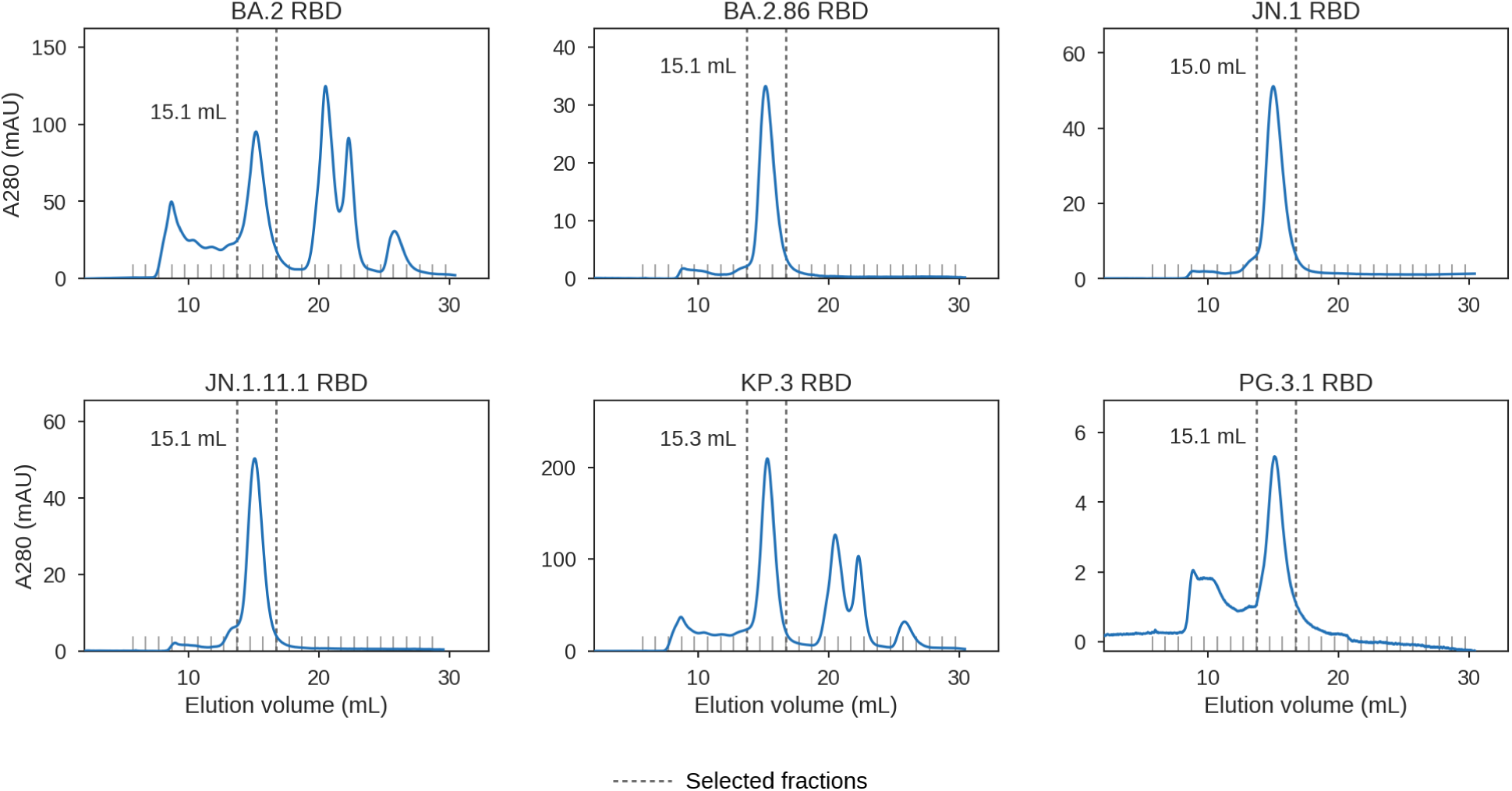
Size-exclusion chromatography of the BA.2-derived RBD variant antigen panel. Size-exclusion chromatograms for BA.2, BA.2.86, JN.1, JN.1.11.1, KP.3 and PG.3.1 RBDs. Traces show absorbance at 280 nm as a function of elution volume. Gray baseline ticks indicate collected fraction boundaries at 1 mL intervals; dashed vertical lines delimit the fractions pooled for downstream experiments (fractions 10–12, 13.7–16.7 mL). Annotated values indicate the elution volume of the principal peak within the selected 13–18 mL product region.

**Figure S27.**
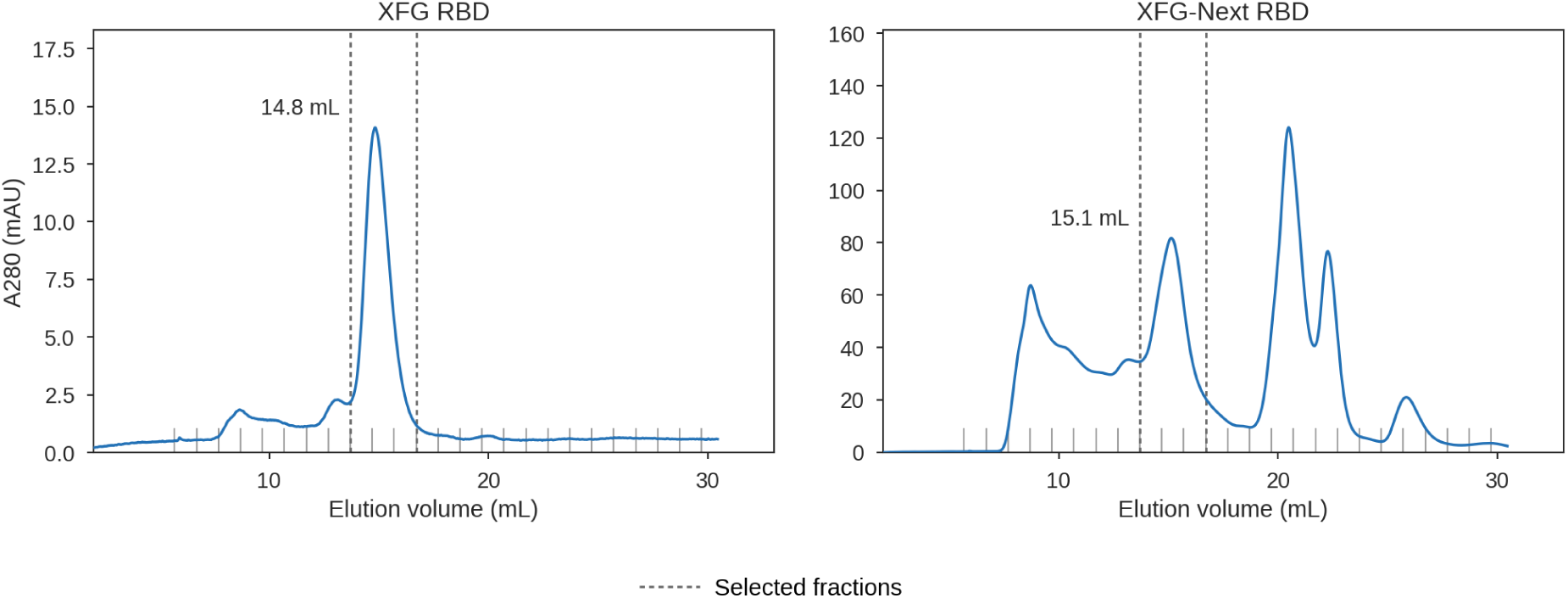
Size-exclusion chromatography of the XFG and XFG-Next RBDs. Size-exclusion chromatograms for the XFG and XFG-Next RBD antigens. Traces show absorbance at 280 nm as a function of elution volume. Gray baseline ticks indicate collected fraction boundaries at 1 mL intervals; dashed vertical lines delimit the fractions pooled for downstream experiments (fractions 10–12, 13.7–16.7 mL). Annotated values indicate the elution volume of the principal peak within the selected 13–18 mL product region.

**Figure S28.**
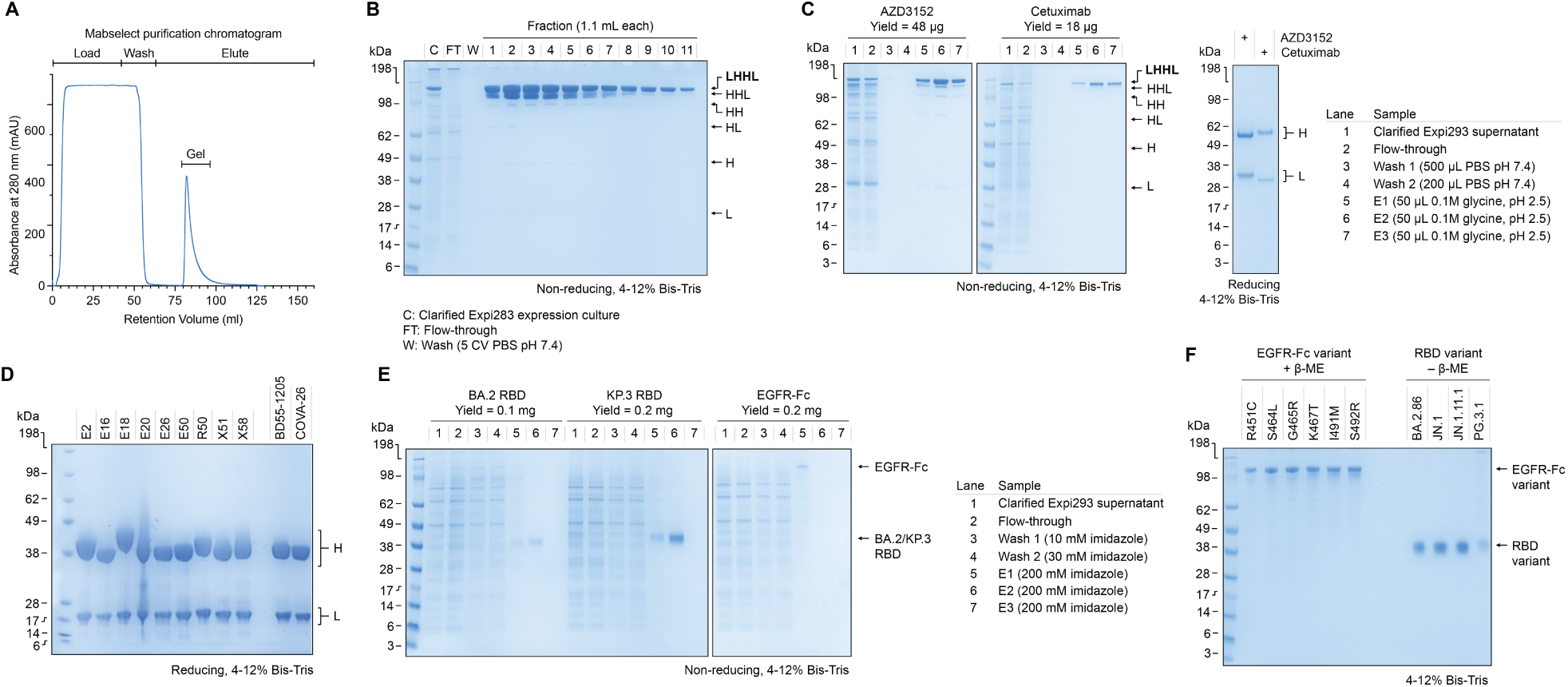
SDS-PAGE characterization of purified antibodies and antigens. **(A)** MabSelect affinity purification chromatogram for AZD3152. **(B)** Non-reducing SDS-PAGE of AZD3152 purification fractions. **(C)** Protein G magnetic bead purification of AZD3152 and cetuximab antibodies, monitored at each step by non-reducing SDS-PAGE. A reducing gel of purified antibodies shows the corresponding heavy (H) and light (L) chains. Total purified yields are indicated above each gel. **(D)** Reducing SDS-PAGE of purified design and control antibody preparations, showing heavy (H) and light (L) chains. **(E)** Immobilized metal affinity chromatography purification of recombinant SARS-CoV-2 BA.2 and KP.3 receptor-binding domains (RBDs) and EGFR-Fc, monitored by non-reducing SDS-PAGE. Total purified yields are indicated above each gel. **(F)** SDS-PAGE of purified EGFR-Fc resistance variants under reducing conditions (+*β*-mercaptoethanol) and SARS-CoV-2 RBD intermediate variants under non-reducing conditions (−*β*-mercaptoethanol). Molecular weights (kDa) are indicated alongside each gel. All gels were run using 4–12% Bis-Tris gels.

**Figure S29.**
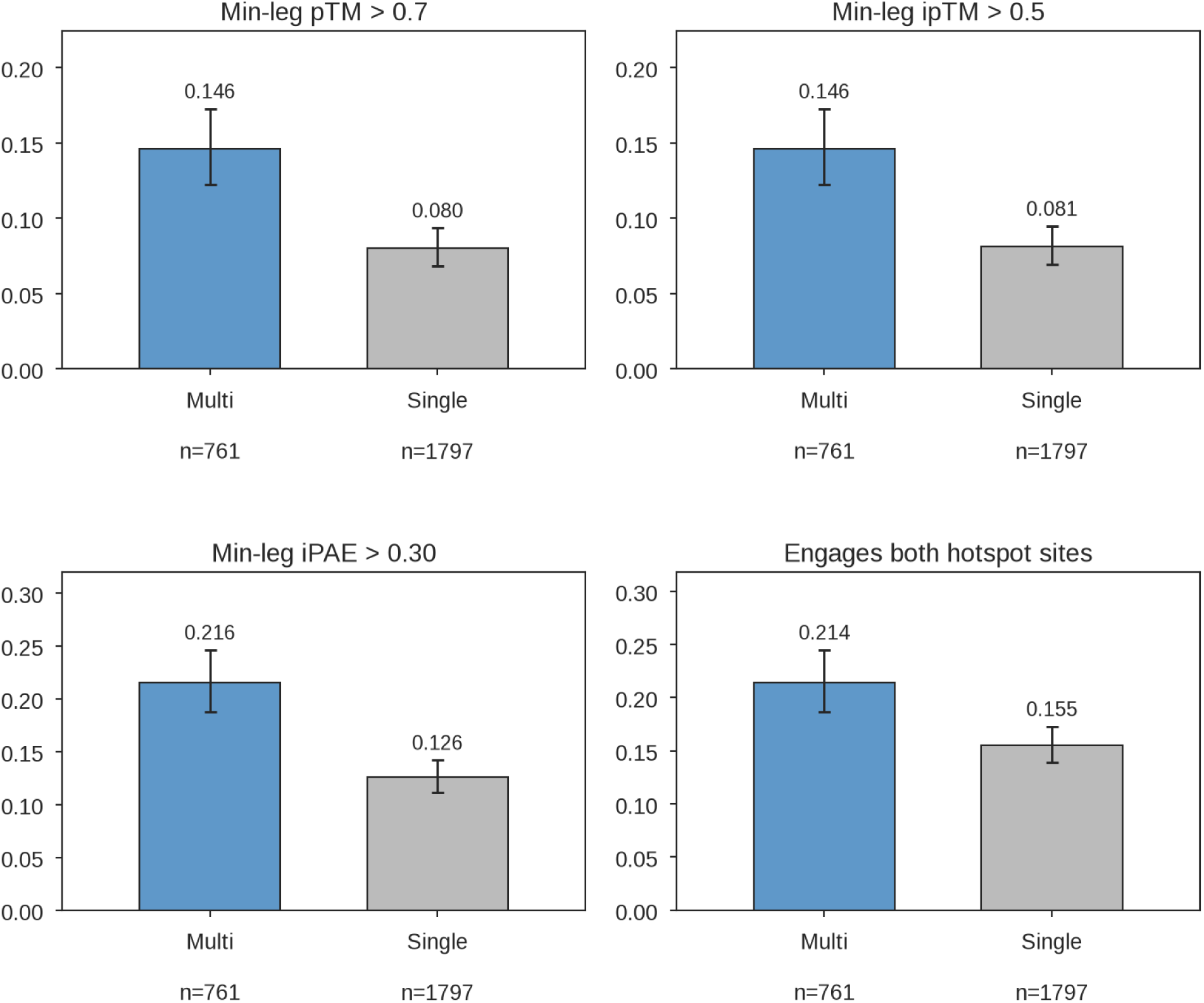
Multi-target binding pass rates for Germinal and Germinal Multi. Designs were generated by co-optimizing against both targets in a campaign (*Multi*, blue) or optimizing against a single target (*Single*, grey). All designs were refolded and scored against both target using AF3, with the worst value across targets used for each metric. Bars pool designs from three campaigns: EGFR clinical escape, BA.2/KP.3, and XFG/XFG-Next. A design engages a hotspot site if at least 75% of the corresponding target’s hotspot residues lie within 4.5 Å of the binder. Interface PAE is reported using pDockQ’s (Bryant et al., 2022) normalized ifpae_norm (0–1, higher is better). Error bars indicate Wilson 95% confidence intervals.

**Figure S30.**
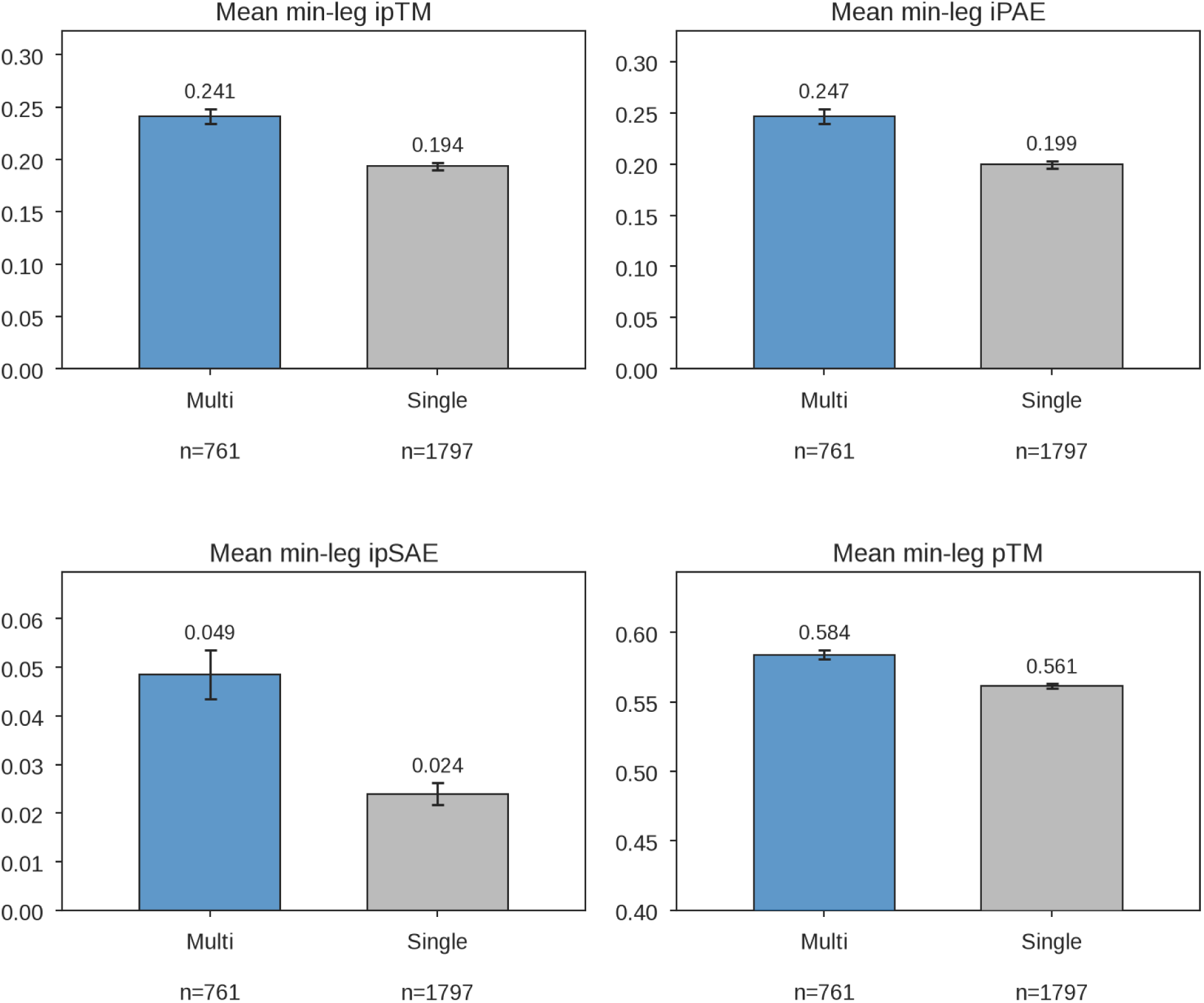
Multi-target binding confidence for Germinal and Germinal Multi. Mean confidence based on the worst value across targets for each design, pooled as in Figure S29. Error bars indicate ±1 SEM.

**Figure S31.**
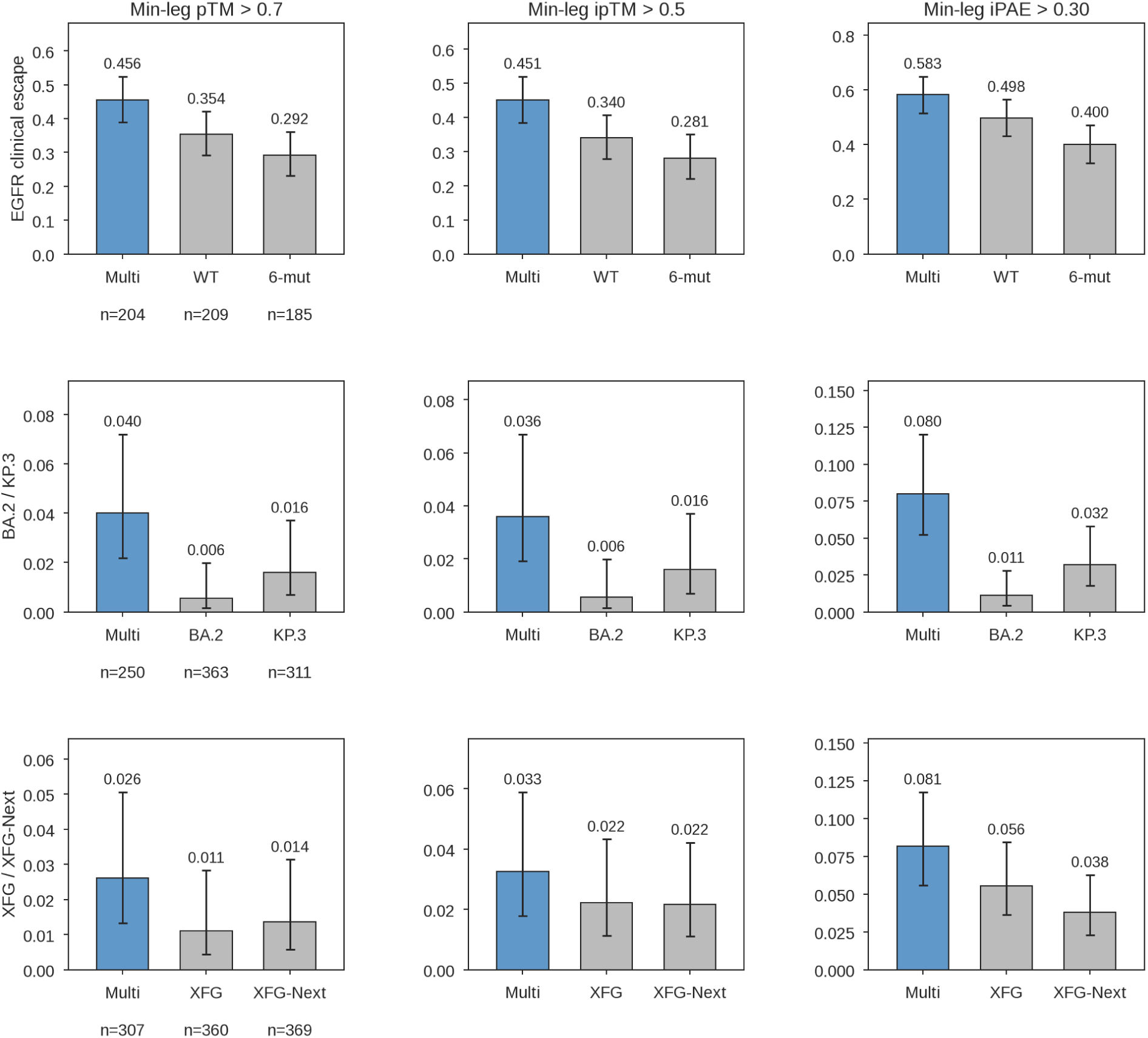
Per-campaign multi-target binding pass rates for Germinal and Germinal Multi. Pass rates from Figure S29, stratified by campaign. Rows represent campaigns, and columns represent the three metrics, with thresholds applied to the worst value across targets. Each panel shows the co-optimized arm (blue) with that campaign’s two single-target arms (grey). Error bars indicate Wilson 95% confidence intervals; sample size (*n*) are reported once per row.

